# Homeostatic Controllers Compensating for Growth and Perturbations

**DOI:** 10.1101/466029

**Authors:** Peter Ruoff, Oleg Agafonov, Daniel M. Tveit, Kristian Thorsen, Tormod Drengstig

## Abstract

Cells and organisms have developed homeostatic mechanisms to maintain internal stabilities which protect them against a changing environment. How cellular growth and homeostasis interact is still not well understood, but of increasing interest to the synthetic and molecular biology community where molecular control circuits are sought and tried to maintain homeostasis that opposes the diluting effects of cell growth. In this paper we describe the performance of four negative feedback (inflow) controllers, which, for different observed growth laws (time-dependent increase in the cellular volume *V*) are able to compensate for various time-dependent removals of the controlled variable *A*. The four implementations of integral control are based on zero-order, first-order autocatalytic, second-order (antithetic), and derepressing inhibition kinetics. All controllers behave ideal in the sense that they for step-wise perturbations in *V* and *A* are able to drive the controlled variable precisely back to the controller’s theoretical set-point
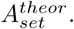
The applied increase in cellular volume includes linear, exponential and saturating growth and reflect experimentally observed growth laws of single cell organisms and other cell types. During the increase in *V*, additional linear or exponential time-dependent perturbations which remove *A* are applied, and controllers are tested with respect to their ability to compensate for both the increase in volume *V* and the applied perturbations removing *A*. Our results show that the way how integral control is kinetically implemented and the structure of the negative feedback loop are essential determinants of a controller’s performance. The results provide a ranking between the four tested controller types. Considering exponential volume increases together with an exponentially increasing removal rate of *A* controllers based on derepression kinetics perform best, but break down when the control-inhibitor’s concentration gets too low. The first-order autocatalytic controller is able to defend time-dependent exponential growth and removals in *A*, but generally with a certain offset below its theoretical set-point
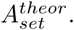
The controllers based on zero-order and second-order (antithetic) integral feedback can only manage linear increases in *V* and removals in *A*, in dependence of the controllers’ aggressiveness. Our results provide a theoretical basis what controller kinetics are needed in order to compensate for different growth laws.

## Introduction

The term *homeostasis* was defined by Walter B. Cannon [1] to describe the coordinated ability of organisms and cells to maintain an internal stability by keeping concentrations of cellular components within certain tolerable limits [2]. Cannon’s emphasis on *homeo* indicates that he considered the internal physiological state not as a constant, as suggested earlier by Benard’s concept of a fixed “milieu intérieur” [2, 3], but conceives homeostasis as a dynamic adaptable system which allows variations within certain limits. Dependent on the controlled components, the homeostatic limits in which one or several controllers operate can vary considerably. For example, while the negative feedback regulation of cellular sodium shows an apparently changing (rheostatic) and less well-defined set-point [4, 5], the regulation of other metal ions including cytosolic calcium have more strict limits [6–8].

Growth, an essential aspect of all living beings is a highly regulated process. Although the protective functions of homeostasis need to be in place during growth, the interacting mechanisms between homeostasis and growth are not well understood. In principle, there are two aspects to consider. The first aspect, which is considered in this paper, is how homeostatic mechanisms can compensate for growth (and additional perturbations) without influencing growth. The other aspect, considered in a following paper, is how homeostatic mechanisms can influence growth. In this paper we consider growth as an increase of the cellular volume. We investigate by testing four negative feedback loop structures/motifs [7] with different kinetic implementations of integral control how these controllers are capable to oppose the dilution effects of growth along with additional perturbations (removals) in a controlled variable. Integral control is a concept from control engineering [9] enabling robust control for step-wise perturbations, and has been implicated to occur in a variety of homeostatic regulated systems [5, 10–12]

The growth kinetics that will be considered include linear (constant) as well as saturating and exponential growth laws. According to Bertalanffy [13, 14], the different observed growth kinetics of organisms are related to the organisms’ metabolism. For example, when respiration is proportional to the surface of the organism linear growth kinetics are obtained. On the other hand, if respiration is proportional to the organism’s weight/volume, exponential growth occurs. Growth kinetics of bacteria [15, 16] appear closely related to the bacterial form or shape. Rod-shaped bacteria show exponential growth rates, i.e.

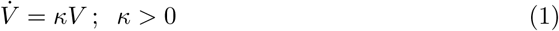

whereas spherical bacteria increase their cellular volume by a rate law related to the
surface to volume ratio, i.e.,

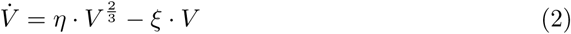

where *η* and *ξ* are constants reflecting anabolism and catabolism, respectively [17].

During growth the controllers have also been tested with respect to outflow perturbations acting on a controlled variable *A*, i.e., by removing *A* in a time-dependent (linear or exponential) manner. We focus here primarily on outflow perturbations, because together with the diluting effects of the different growth laws these perturbations represent the most severe conditions for testing the controllers.

## Materials, methods, and controllers tested

For the sake of simplicity we assume that compounds inside a growing cellular volume *V* undergo ideal mixing. Computations were performed by using the Fortran subroutine LSODE [18]. Plots were generated with gnuplot (www.gnuplot.info) and Adobe Illustrator (adobe.com). To make notations simpler, concentrations of compounds are denoted by compound names without square brackets. Time derivatives are indicated by the ‘dot’ notation. Concentrations and rate parameter values are given in arbitrary units (au). Rate parameters are presented as *k_i_*’s (*i*=1, 2, 3, …) irrespective of their kinetic nature, i.e. whether they represent turnover numbers, Michaelis constants, or inhibition constants. A set of MATLAB (mathworks.com) calculations with instructions are provided in the Supporting Information as a combined zip-file (S1 Matlab). In addition, the Supporting Information contains analytical expressions/estimates of steady state values.

We have previously described eight different two-component negative feedback loop arrangements (motifs) dependent on how the two components *A* (the controlled variable) and *E* (controller variable) activate or inhibit each other [7]. Half of the motifs describe inflow controllers, i.e., the compensatory flux adds *A* to the system from a certain source, while the other half are outflow controllers in which the compensatory flux removes *A* in order to maintain homeostasis in *A*. Since our interest here is to study controller performances that compensate for the dilution effects of growth together with (time-dependent) perturbations which remove *A*, we focus here entirely on inflow controllers. From the four inflow controller motifs we have chosen two, one based on only activation (motif 1, [7]), while the other is based on both activation and inhibition/derepression (motif 2). The reason for this choice comes from an earlier study [19] on time-dependent perturbations which indicated that a motif 1 controller with an autocatalytic (positive feedback) implementation of integral control [6, 9, 20] and a zero-order motif 2 controller with derepression kinetics showed good performances in comparison with a zero-order based motif 1 controller. In addition, we have included a motif 1 based controller with a second-order (antithetic, [21]) implementation of integral control.

Fig. 1 gives an overview of the tested controller types. All controllers behave ideal in the sense that they for step-wise removals in *A*, and in the absence of growth, are able to keep *A* precisely at their defined theoretical set-points
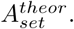
The controllers were investigated with respect to their capabilities to compensate for time-dependent outflow perturbations in *A* and in the presence of different growth laws (increase in the reaction volume *V*) according to Bertalanffy’s classifications [13, 14, 17].

**Figure 1.**
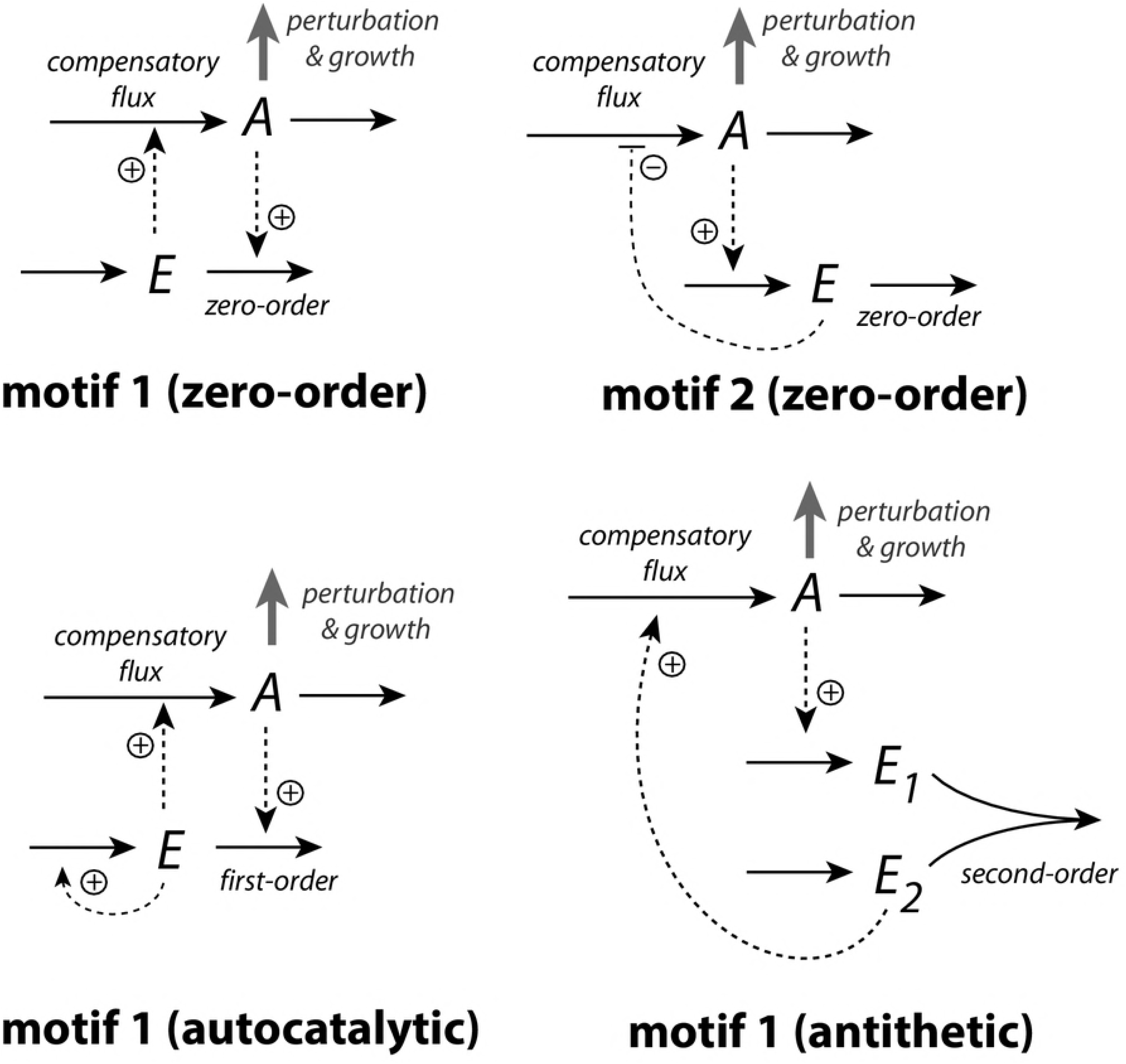
The controllers investigated in this study. Reaction orders are with respect to *E*. The reaction between *E*_1_ and *E*_2_ in the antithetic controller is an overall second-order process.

## Reaction kinetics during volume changes

To get a correct description of the cellular concentration changes during cell growth we have to consider the concentration changes due to the increasing reaction volume *V*. If *A* denotes the concentration of *n_A_* moles of compound *A* in volume *V*, the overall change of concentration *A* is composed of two terms, one that describes the changes of *A* while *V* is kept constant, *Ȧ_V_* and of a second term, *A*(*V̇/V*), which describes the influence of the volume changes on the concentration of *A*, i.e.,

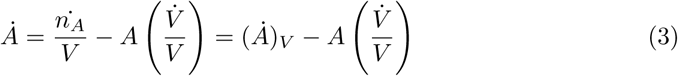

The structure of Eq. 3 will be used when formulating the rate equations of cellular compounds in the presence of a changing *V*. Before we turn to the actual controller examples we show how growth (*V̇*) affects the concentration of a given species *A* (which will be later our controlled variable) when *A* is unreactive, being produced internally within the cell, or being produced by a transporter-mediated process.

### Unreactive *A*

In this example (Fig. 2) *n_A_* is kept constant, but the volume *V* increases with the rate *V̇*.

**Figure 2.**
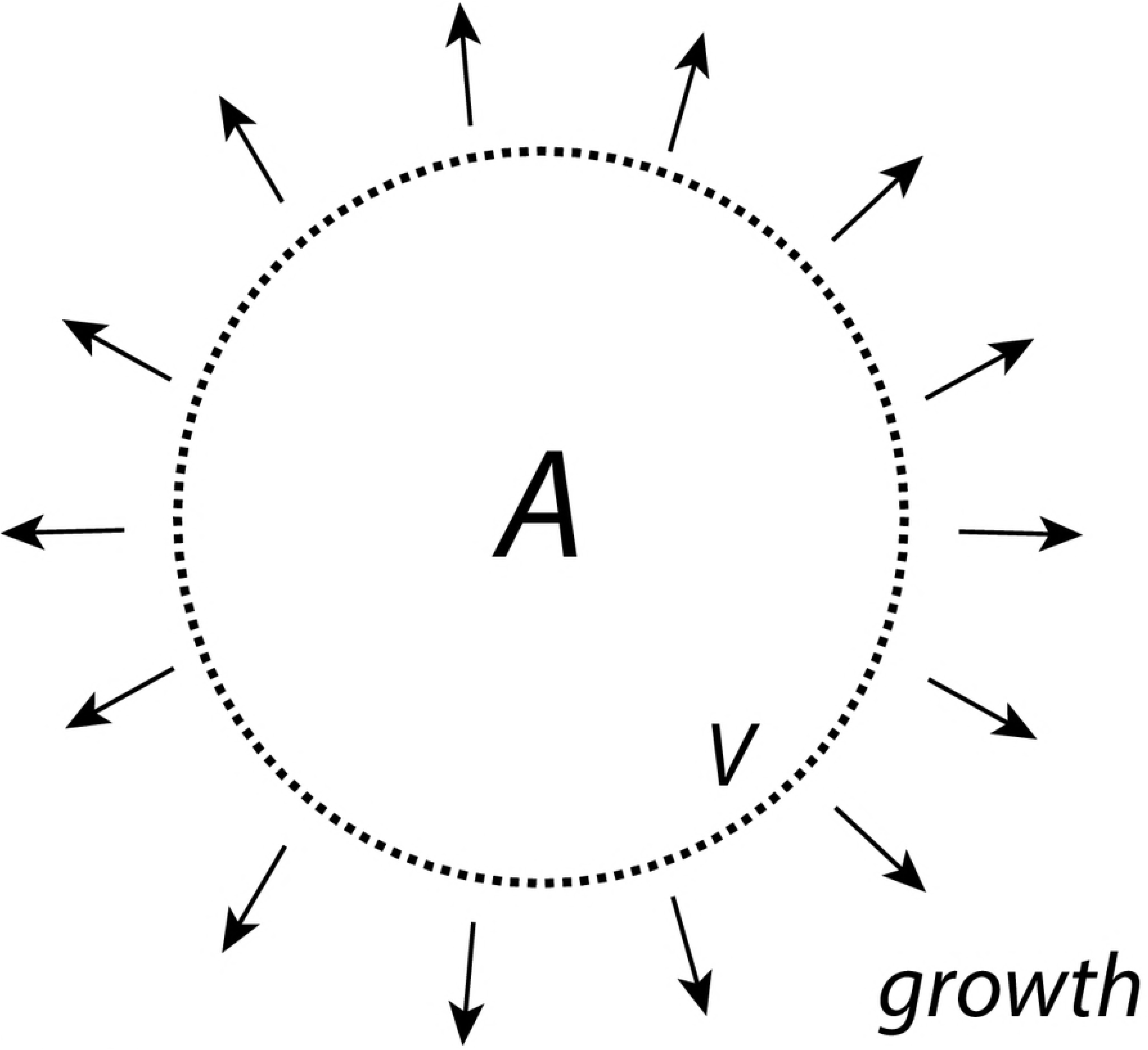
*A* is present inside the cell with a constant amount of *n_A_* moles, while the cellular volume *V* increases with rate *V̇*.

As *V* increases the concentration of *A* will decrease, i.e.,

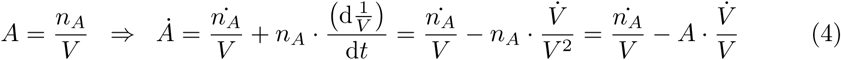

Since we assume that *n_A_* is constant, we have that
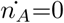
and the concentration of *A* decreases according to

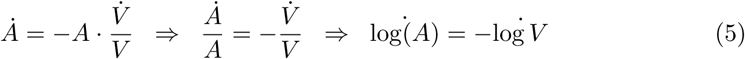

Integrating Eq. 5 leads to:

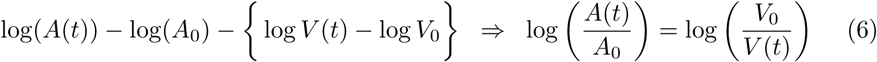

which can be rewritten as

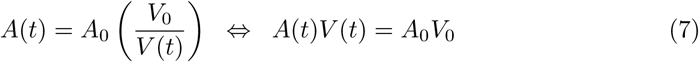

Eq. 7 can also be derived by noting that *A*_0_=*n_A_/V*_0_ and *A*(*t*)=*n_A_/V* (*t*). Solving for *n_A_* from one of the equations and inserting it into the other leads to Eq. 7.

### Cell internal generated *A*

Compensatory fluxes to counteract diminishing levels of a controlled compound *A* can be generated by a cell internal compound (assumed here to be homogeneously distributed inside *V*) or by the help of transporters from stores outside of the cell or from cell-internal (organelle) stores. We will investigate both ways to generate compensatory fluxes.

To achieve a constant level of *A* from a cell internal source, despite increasing *V*, we consider first a zero-order enzymatic reaction where enzyme *E* converts a species *S* (assumed to be present in sufficiently high amounts) to *A*, where *V* is assumed to increase by a constant rate (Fig. 3).

**Figure 3.**
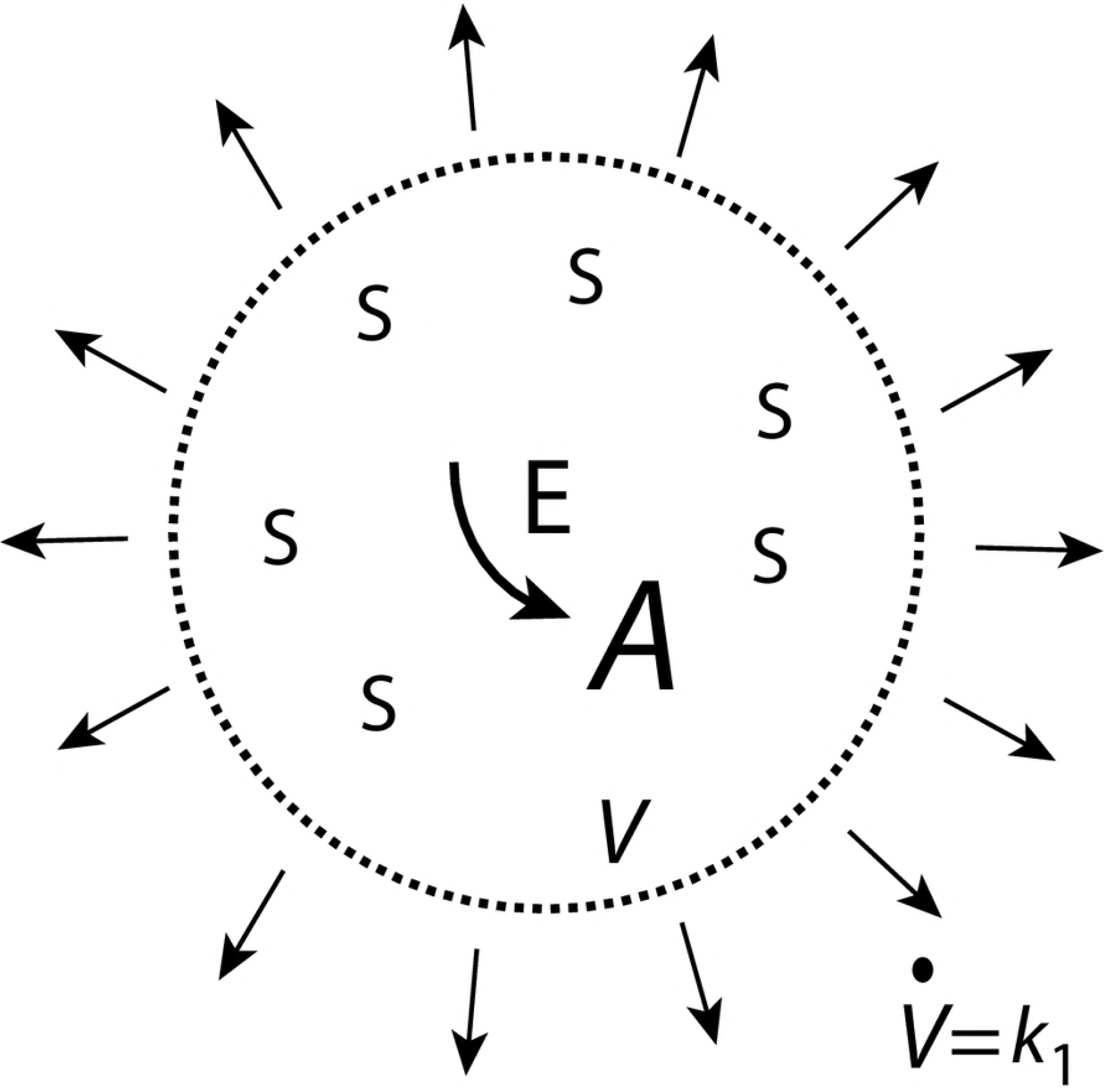
*A* is formed by zero-order kinetics within the cell while the cellular volume increases with a constant rate *V̇* = *k*_1_.

Let’s also assume that *E* is not subject to any synthesis, but that during the increase of *V*, *E* remains always saturated with *S* and produces *A* by zero-order kinetics with respect to *A*. The initial production rate of *A* at time *t*=0 is given as

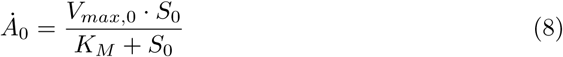

Since *E* is considered to be saturated by *S* at all times we have that *K_M_* ≪ *S*(*t*) leading to

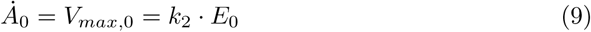

where *k*_2_ is the turnover number of the enzymatic process generating *A*, and *E*_0_ is the enzyme concentration at time *t* = 0. As *V* increases, the concentrations of *E* and *A* are subject to dilution as described by the rate equations

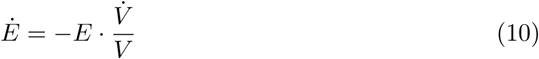

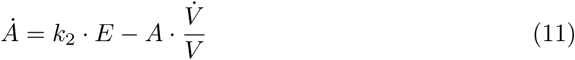

For *V̇* = *k*_1_=constant, *E*(*t*) and *A*(*t*) are described by the equations (S1 Text)

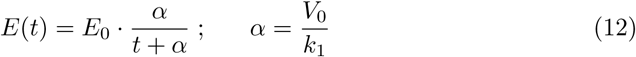

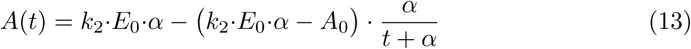

From Eq. 13 we see that *A* will approach a final concentration *A*_final_ = *k*_2_·*E*_0_·*α* even when *V* continues to grow. The time needed of *A* to approach *A*_final_ is determined by the term *α/*(*t*+*α*).

Fig. 4 shows that *A*_final_ is independent of the initial values of *A*. However, the system is not stable against perturbations which remove *A*. In such a case *A* will go to zero (S1 Text).

**Figure 4.**
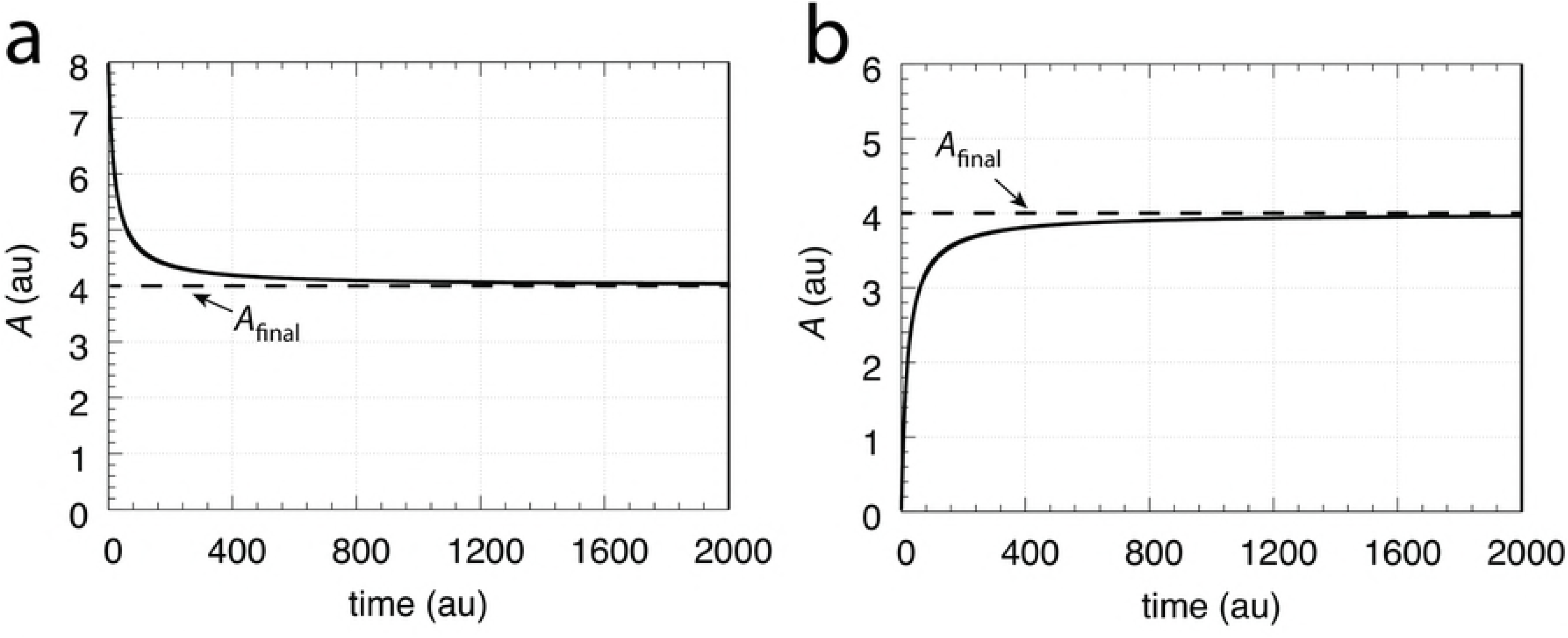
*A* approaches *A*_final_ independent of the initial concentration of *A*. (a) *A*_0_=8.0; (b) *A*_0_=0.0. All other rate parameters are: *k*_1_=*V̇* =1.0, *k*_2_=2.0,*E*_0_=0.1, *V*_0_=20.0.

### Transporter generated *A*

Alternatively, *A* may be imported into the cell by a transporter *T* (Fig. 5).

**Figure 5.**
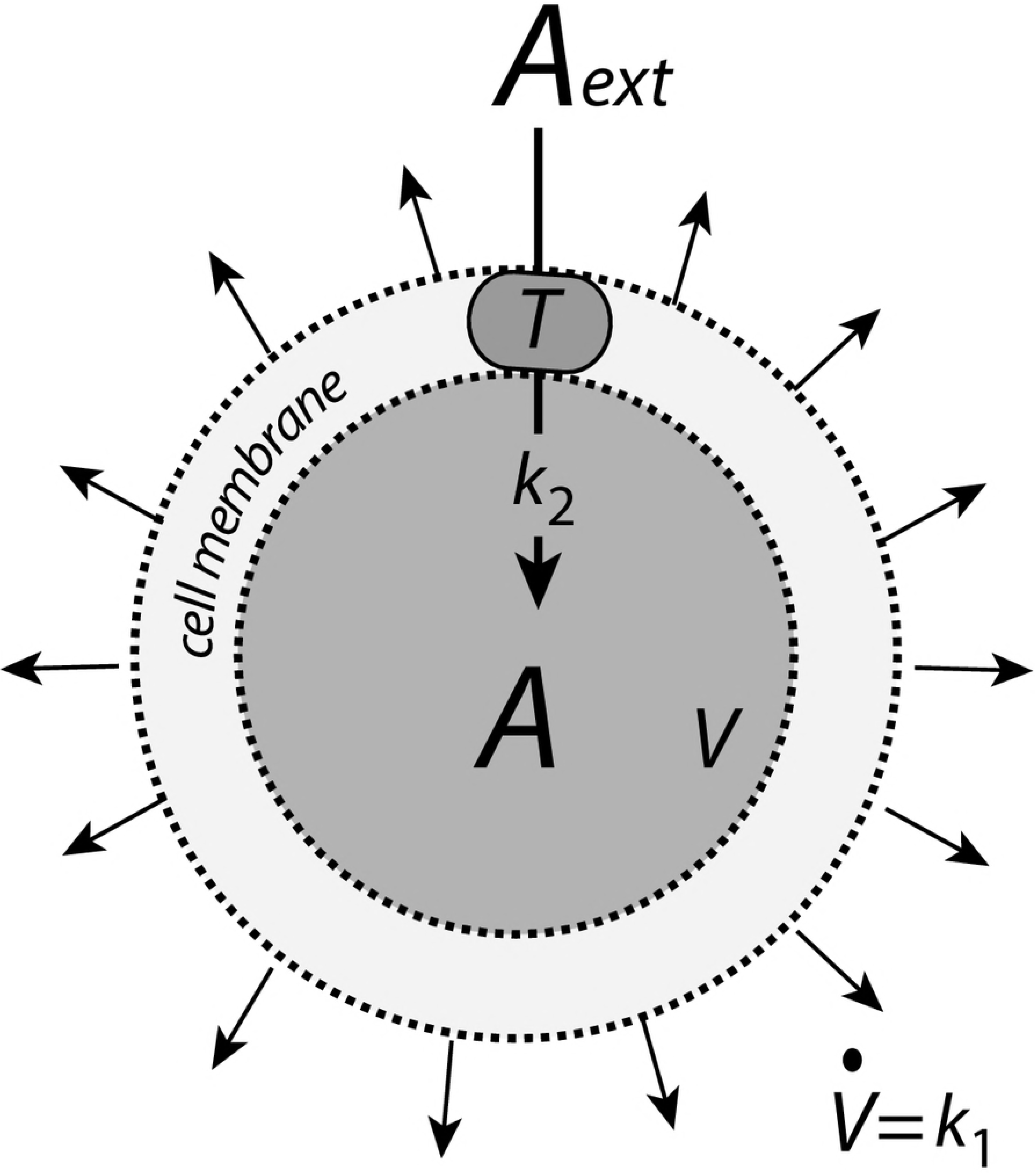
*A* is imported into the cell by transporter *T*.

Also here we consider that the transporter works under saturation (zero-order) conditions adding
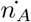
moles of *A* per time unit into the cellular volume *V*

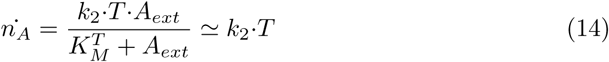

where *T* denotes the (surface/membrane) concentration of the transporter,
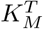
is a dissociation constant between external *A* (*A_ext_*) and *T*, and *k*_2_ is the turnover number of the transporter-mediated uptake of *A*.

The change in the concentration of *A* inside an expanding cell is given by (see Eq. 3)

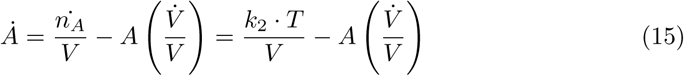

For constant *V̇*, *k*_2_, and *T* the steady state of *A* (*Ȧ*=0) is *k*_2_*T/V̇* independent of the initial concentration of *A*. However, also in the transporter-based inflow of *A*, the steady state in *A* is not stable against perturbations removing *A*. Any reaction within the cell removing *A* while growth occurs will drive *A* to zero (S2 Text). To get a steady state that is stable against perturbations a negative feedback controller needs to be included.

## Controllers with transporter-based compensatory fluxes and linear time-dependent perturbations

In this section the four controller motifs (Fig. 1) use transporter-based compensatory fluxes. The controllers’ performances are tested with respect to constant growth *V̇* =*k*_1_ and a linear increase in the outflow perturbation rate parameter *k*_3_.

### Motif 1 zero-order controller

Fig. 6 shows the motif 1 controller with zero-order implementation of integral control [7]. *A* is the controlled compound and *E* is the controller molecule which concentration (in the ideal controller case) is proportional to the integrated error between *A* and
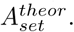
*M* i a precursor/store or *E*, which when consumed limits the operational life time of the controller.

**Figure 6.**
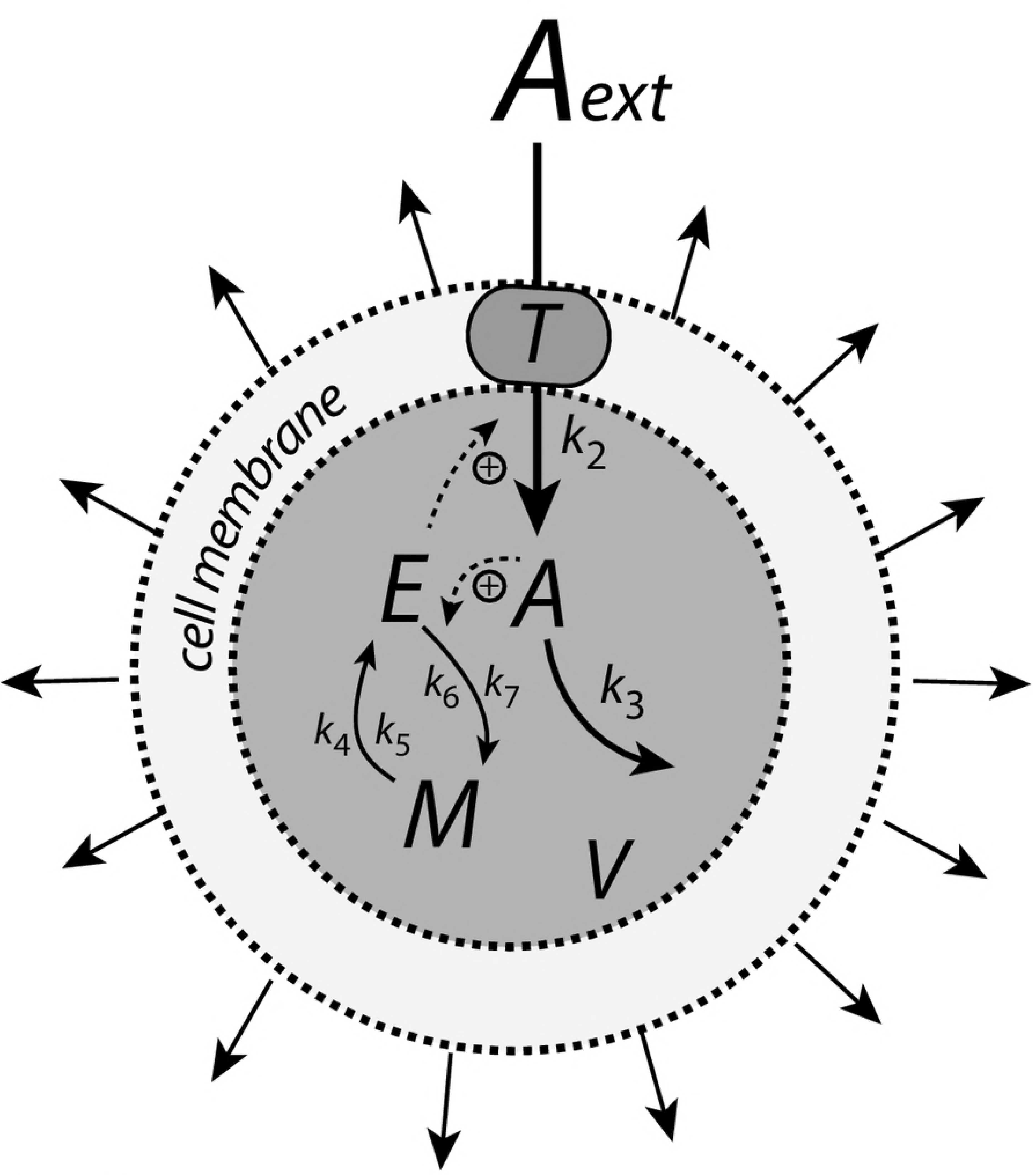
Controller based on motif 1 zero-order integral control with transporter generated compensatory flux. The controller species *E* is produced by an enzymatic zero-order process from compound *M*. *E* is recycled by another zero-order process (with respect to *E*) but the rate of *E*-removal is proportionally to the concentration of *A*. Outflow perturbations are represented by the rate *r*_3_=*k*_3_·*A*, where *k*_3_ is either constant or increases linearly with time.

The rate equations for this system are:

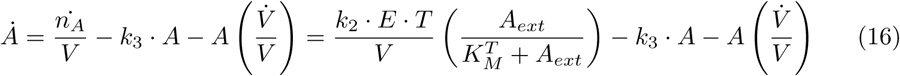

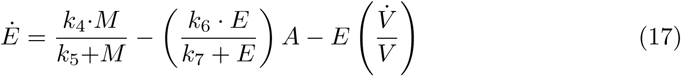

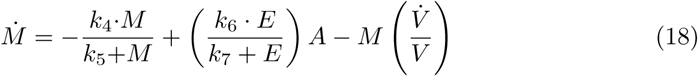

For simplicity *T* and *A_ext_*/(
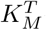
+ *A_ext_*) are set to 1 leading to an inflow rate in *A* of *k*_2_*E/V*. When *k̇*_3_=*V̇* =0, the set-point of the controller is (Ref. [7], S3 Text)

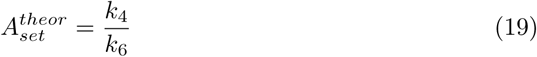

independent of the inflow rate constant *k*_2_ and the perturbation strength parameter *k*_3_.

When *V̇* =constant the zero-order controller maintains a steady state below
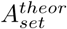
(S3 Text):

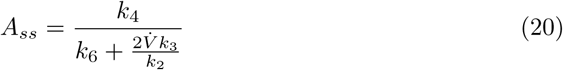

which becomes dependent of *V̇*, and the rate constants *k*_2_ and *k*_3_.

In testing the performance of this controller we consider three phases (see Fig. 7). During the first phase the volume and the perturbation *k*_3_ are kept constant. The controller is able to compensate for the perturbation rate *k*_3_·*A* and keeps *A* at its theoretical set-point
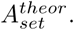
In the second phase the volume increases linearly, while *k*_3_ remains constant. The zero-order controller is now no longer able to maintain homeostasis at
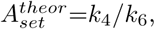
but shows a *V̇*-dependent offset below
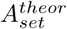
as described by Eq. 20. When in phase 3 also *k*_3_ increases linearly the controller breaks down and *A* goes to zero.

**Figure 7.**
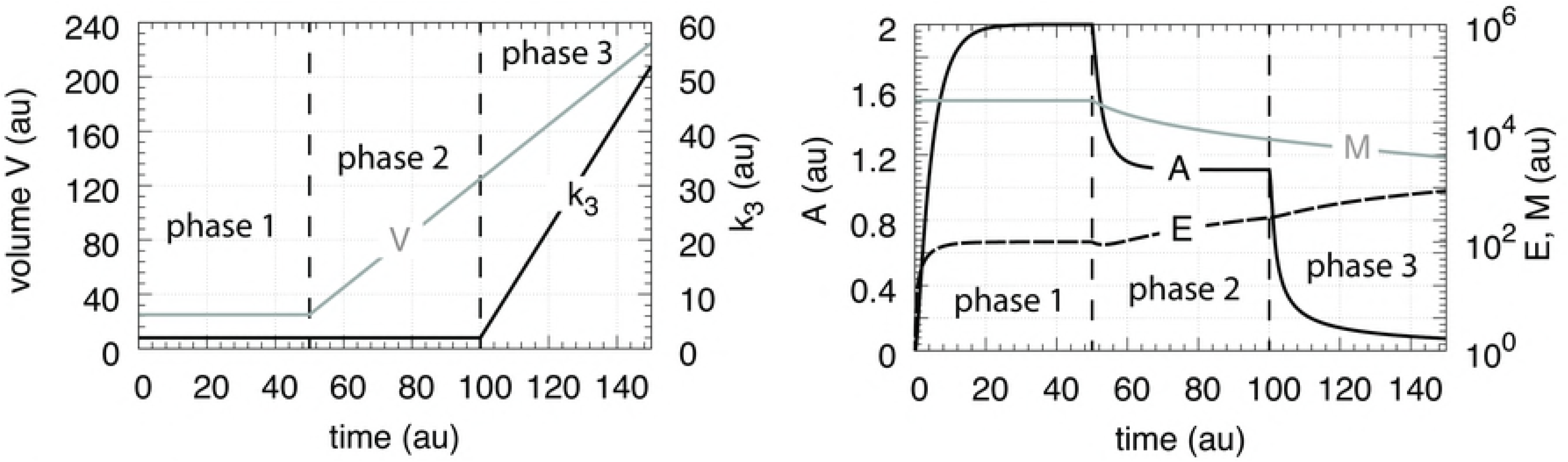
Performance of the motif 1 zero-order controller with transporter mediated compensatory flux (Eqs. 16-18). Phase 1: constant volume *V* and constant *k*_3_. Initial concentrations and rate constant values: *V*_0_=25.0, *V̇* =0.0, *A*_0_=0.0, *E*_0_=0.0, *M*_0_=4 × 10^4^, *k*_2_=1.0, *k*_3_=2.0, *k̇*_3_=0.0, *k*_4_=20.0, *k*_5_=1 × 10^−6^, *k*_6_=10.0, *k*_7_=1 × 10^−6^. The controller keeps *A* to its theoretical set-point at
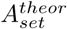
=*k*_4_/*k*_6_=2.0 (Eq. 19). Phase 2: rate constants remain the same as in phase 1, but *V* increases now linearly with *V̇* =2.0, while *k*_3_ remains constant at *k*_3_=2.0. The controller shows an offset below
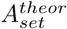
with *A_ss_*=1.11 in agreement with Eq. 20. Phase 3: *V* continues to increase with the same speed while *k*_3_ now linearly increases with *k̇*_3_=1.0. As indicated by Eq. 20 the controller breaks down, *A* goes to zero, as *V̇* and *k*_3_ increase.

### Motif 1 antithetic controller

The antithetic controller [21] uses two controller molecules, *E*_1_ and *E*_2_ (Fig. 8). Compound *E*_1_ is activated by *A* but is removed by compound *E*_2_ by a second-order process. *E*_2_ is formed by a zero-order process which acts as a constant reference rate. In addition, *E*_2_ also acts as a signaling molecule, which closes the negative feedback loop by activating the transporter-based compensatory inflow of *A*.

**Figure 8.**
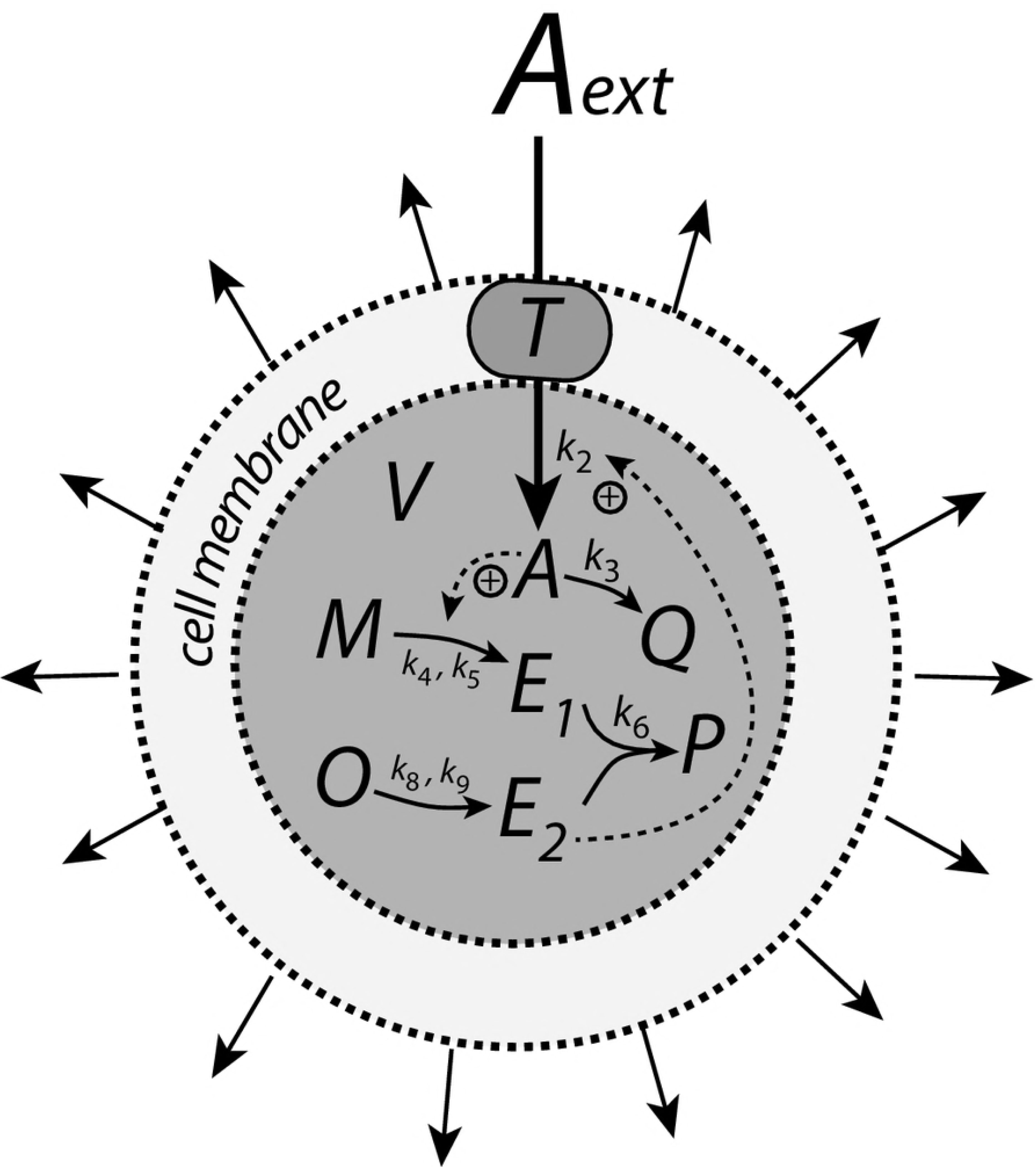
Motif 1 based controller with second-order (antithetic) integral control. The controller species *E*_2_ is produced by an enzymatic zero-order process from compound *O*. *E*_2_ activates the transporter-based compensatory flux of *A* and is removed by *E*_1_ using second-order kinetics forming *P*.

Assuming, as in the previous two examples that *T* and *A_ext_/*(
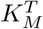
+*A_ext_*) are both 1, the rate equations are

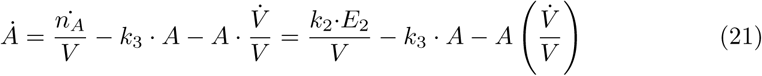

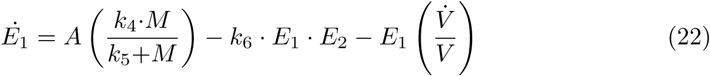

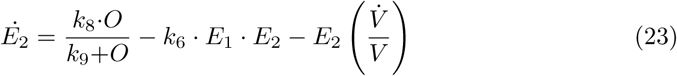

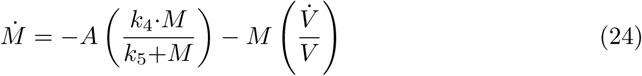

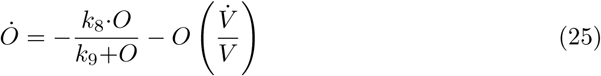

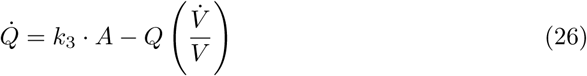

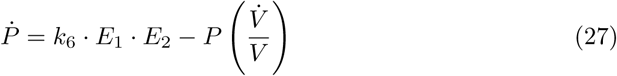

where *k*_5_ ≪ *M* and *k*_9_ ≪ *O* such that the generation of *E*_1_ and *E*_2_ are zero-order processes with respect to *M* and *O*.

In case *V̇* =0 and *k̇*_3_=0 the set-point of the controller is given by setting Eq. 22 and Eq. 23 to zero. Eliminating the second-order term *k*_6_·*E*_1_·*E*_2_ leads to

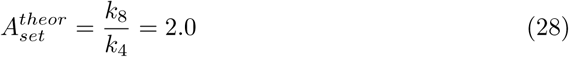

which is shown in phase 1 of Fig. 9. In phase 2 the volume increases linearly with *V̇* =2.0 (Fig. 9, left panel) while *k*_3_ remains to be constant at *k*_3_=2.0. The controller is no longer be able to keep *A* at its theoretical set-point (Eq. 28). When *V̇* and *k*_3_ are constant an analytical expression of *A_ss_* can be derived in good agreement with the numerical calculations (S4 Text):

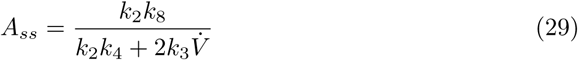

which is analogous to the *A_ss_* expression of the motif 1 zero-order controller (Eq. 20). Finally, in phase 3 *k*_3_ increases linearly with *k̇*_3_=1 together with *V̇* =2.0. As indicated by Eq. 29 and shown by the numerical calculations (Fig. 9) the antithetic controller, like the zero-order controller, breaks down and *A* goes to zero (S4 Text).

**Figure 9.**
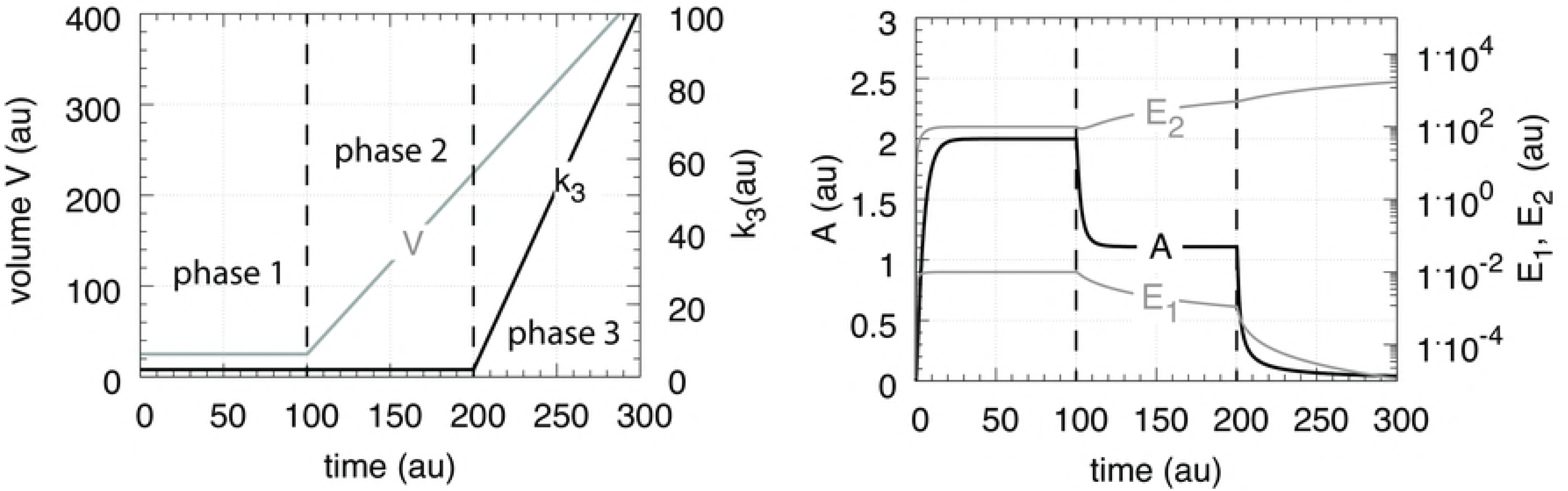
Performance of the antithetic controller with transporter mediated compensatory flux (Eqs. 21-27). Phase 1: constant volume *V* and constant *k*_3_. Initial concentrations and rate constant values: *V*_0_=25.0, *V̇* =0.0, *A*_0_=0.0, *E*_1,0_=0.0, *E*_2,0_=0.0, *M*_0_=1 × 10^5^, *O*_0_=1 × 10^5^, *k*_2_=1.0, *k*_3_=2.0, *k̇*_3_=0.0, *k*_4_=10.0, *k*_5_=1 × 10^−6^, *k*_6_=20.0, *k*_7_ not used, *k*_8_=20.0, *k*_9_=1 × 10^−6^. The controller keeps *A* at its theoretical set-point at
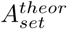
=*k*_8_/*k*_4_=2.0 (Eq. 28). Phase 2: rate constants remain the same as in phase 1, but
*V* increases linearly with *V̇* =2.0, while *k*_3_ remains constant at *k*_3_=2.0. The controller shows an offset below
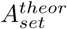
with *A_ss_*=1.11 in agreement with Eq. 29. Phase 3: *V* continues to increase while *k*_3_ increases linearly with *k̇*_3_=1.0. As indicated by Eq. 29 the controller breaks down and *A* goes to zero.

Although not shown explicitly here, the following mass balances are obeyed:

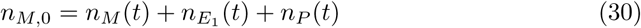

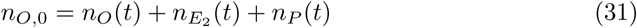

where *n*_*i*,0_ and *n_i_* are respectively the initial number of moles and the number of moles at time *t* of compound *i*.

As described above, when using a transporter mediated compensation in *A* the antithetic and the motif 1 zero-order controller have to increase their controller variables *E*_2_ or *E* in order to keep *A_ss_* constant, as indicated by the equation

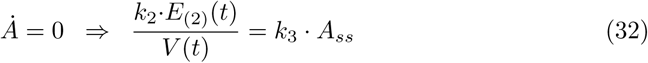

where *E*_(2)_ represents *E*_2_ or *E*.

### Motif 1 autocatalytic controller

Similar to controllers based on double integral action [22] an autocatalytic design [20] is able to keep the controlled species at its set-point even when perturbations become linearly time dependent and rapid [19]. However, in contrast to double integral action the autocatalytic controller is able to compensate for time-dependent perturbations of the form *a*·*t^n^* where *n* is larger than 1.

Fig. 10 shows the reaction scheme. The controller compound *E* is produced autocatalytically, i.e., its rate is proportionally to the concentration of *E*, while *M*, present in relative large amounts, produces *E* by an enzyme-catalyzed reaction which is zero-order with respect to *M*. *E* increases the activity of transporter *T* and leads to an increased import of external *A* into the cell. The negative feedback is closed by an *A*-induced recycling of *E* to *M*. Rate constant *k*_3_ represents a variable perturbation which removes *A* by a first-order process with respect to *A*. The rate equations are:

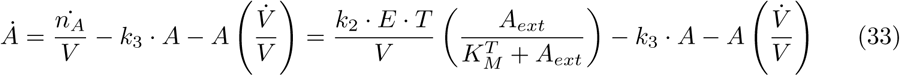

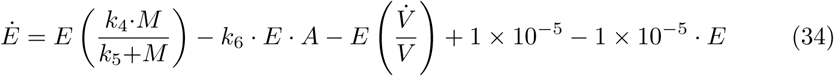

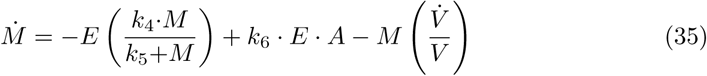

**Figure 10.**
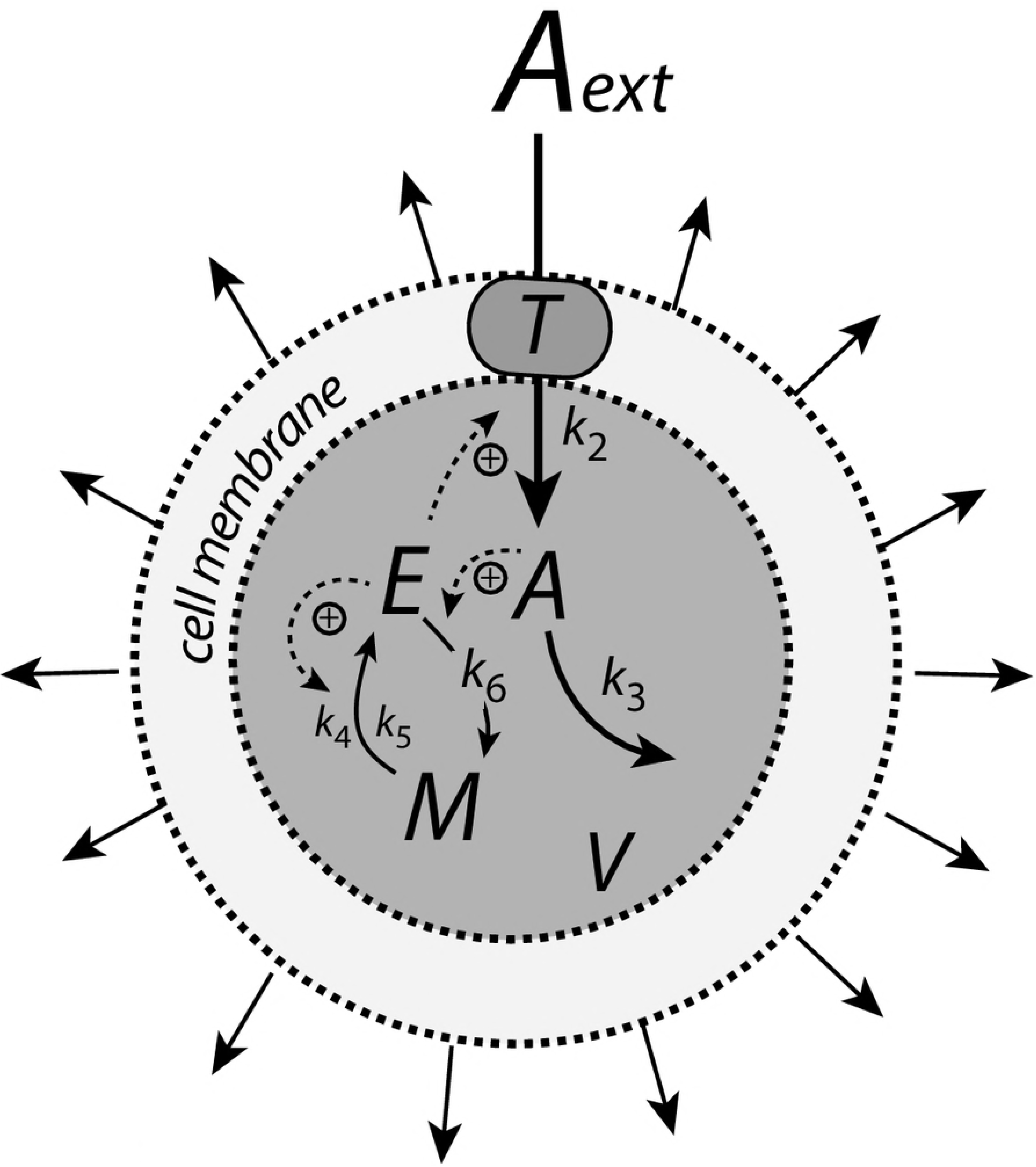
Controller design based on motif 1 autocatalytic integral control. The controller species *E* is produced by an enzymatic zero-order process from compound *M*, but *E* activates its own production and the transporter-based compensatory flux. The negative feedback is due to an inflow activation of *A* by *E* through transporter *T*, while *A* activates the recycling of *E* to *M*. Outflow perturbation in *A* is described by the rate *k*_3_·*A*, where *k*_3_ is either a constant or increases linearly with time.

For simplicity, in Eq. 33, *T* and *A_ext_/*(
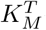
+*A_ext_*) are set to 1. The last two terms in Eq. 34, 1 × 10^−5^ − 1 × 10^−5^ · *E*, represent small background reactions keeping *E* at a sufficiently high level such that the autocatalysis in *E* can start out from zero initial *E* concentrations (see also Ref. [19]).

To determine the controller’s set-point at constant *V* and *k*_3_ we set Eq. 34 to zero. Neglecting the 1 × 10^−5^ − 1 × 10^−5^ · *E* term and setting *V̇* =0, we can solve for the steady state value of *A*, which defines the controller’s theoretical set-point
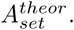

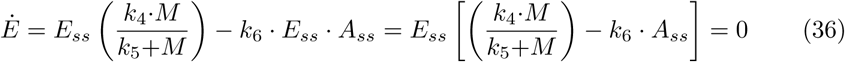

Since *M/*(*k*_5_+*M*)=1 (ideal zero-order conditions), we get from Eq. 36

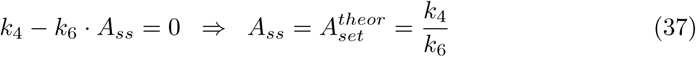

In the case both *V̇* and *k̇*_3_ are nonzero and constant, an (analytical) estimation of the steady state in *A* requires to consider *Ä* (”acceleration” in *A*). For constant *V̇* and *k̇*_3_ values the set-point is calculated to be (S5 Text)

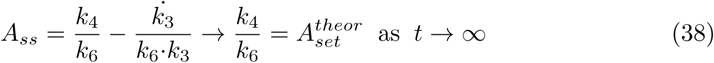

According to previous findings on the autocatalytic controller [19], any time-dependent function *k*_3_(*t*) = *k*_3,0_ + *a*·*t^n^* where *a, n>*0 will lead to the set-point conditions described by Eq. 38 (S5 Text).

The recycling scheme between *E* and *M* implies that *E* and *M* obey a mass balance of the form

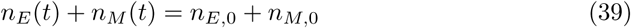

with *n_E_* (*t*)=*E*(*t*)·*V* (*t*), *n_M_* (*t*)=*M* (*t*)·*V* (*t*), and where *n*_*E*,0_ and *n*_*M*,0_ are the initial number of moles of respectively *E* and *M*. The rates how *n_E_* and *n_M_* change at a given time *t* are given as (S5 Text)

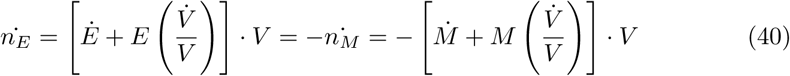

Fig. 11 shows the results. During the first phase no volume change occurs and *k*_3_ is a constant. The controller keeps *A* at
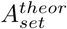
=2.0 as described by Eq. 37. During the second phase both *V* and *k*_3_ increase linearly and the controller still keeps *A* at
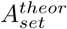
=2.0 according to Eq. 38. To keep *A* at its set-point during increasing *V* and/or *k*_3_ the concentration of *E* has to increase in order to maintain the steady state condition given by Eq. 33 when *Ȧ* =0 and *V̇/V* → 0, i.e.,

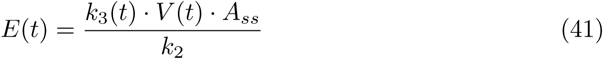

**Figure 11.**
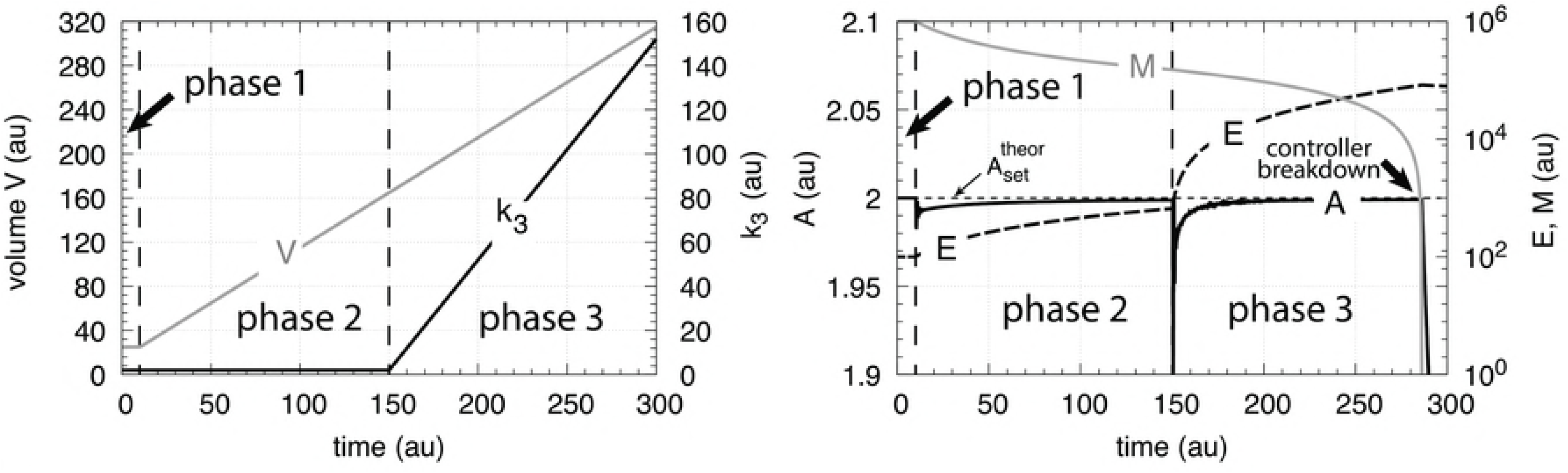
Performance of the motif 1 autocatalytic controller (Eqs. 33-35). Phase 1: constant volume *V* and constant *k*_3_. Initial concentrations and rate constant values: *V*_0_=25.0, *V̇* =0.0, *A*_0_=2.0, *E*_0_=100.0, *M*_0_=1 × 106, *k*_2_=1.0, *k*_3_=2.0, *k̇*_3_=0.0, *k*_4_=20.0, *k*_5_=1 × 10^−6^, *k*_6_=10.0. The controller moves *A* to its set-point at
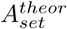
=*k*_4_/*k*_6_=2.0. Phase 2: rate constants remain the same as in phase 1, but *V* increases linearly with *V̇* =1.0. Phase 3: *V* continues to decrease with the same rate and *k*_3_ increases with rate *k̇*_3_=1.0. The controller moves *A* towards
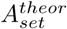
in both phase 2 and phase 3, but breaks down when no additional *E* becomes available by *M* (indicated by the arrow in the right panel).

From the initial conditions (see legend of Fig. 11) we have that *n_E_* (*t*)+*n_M_* (*t*) = *V*_0_ · *M*_0_=2.5 × 10^7^.

### Motif 2 zero-order controller

The reaction scheme of this controller is shown in Fig. 12. The transporter-based compensatory flux is regulated by *E* through repression or derepression and *E* is removed by a zero-order reaction.

**Figure 12.**
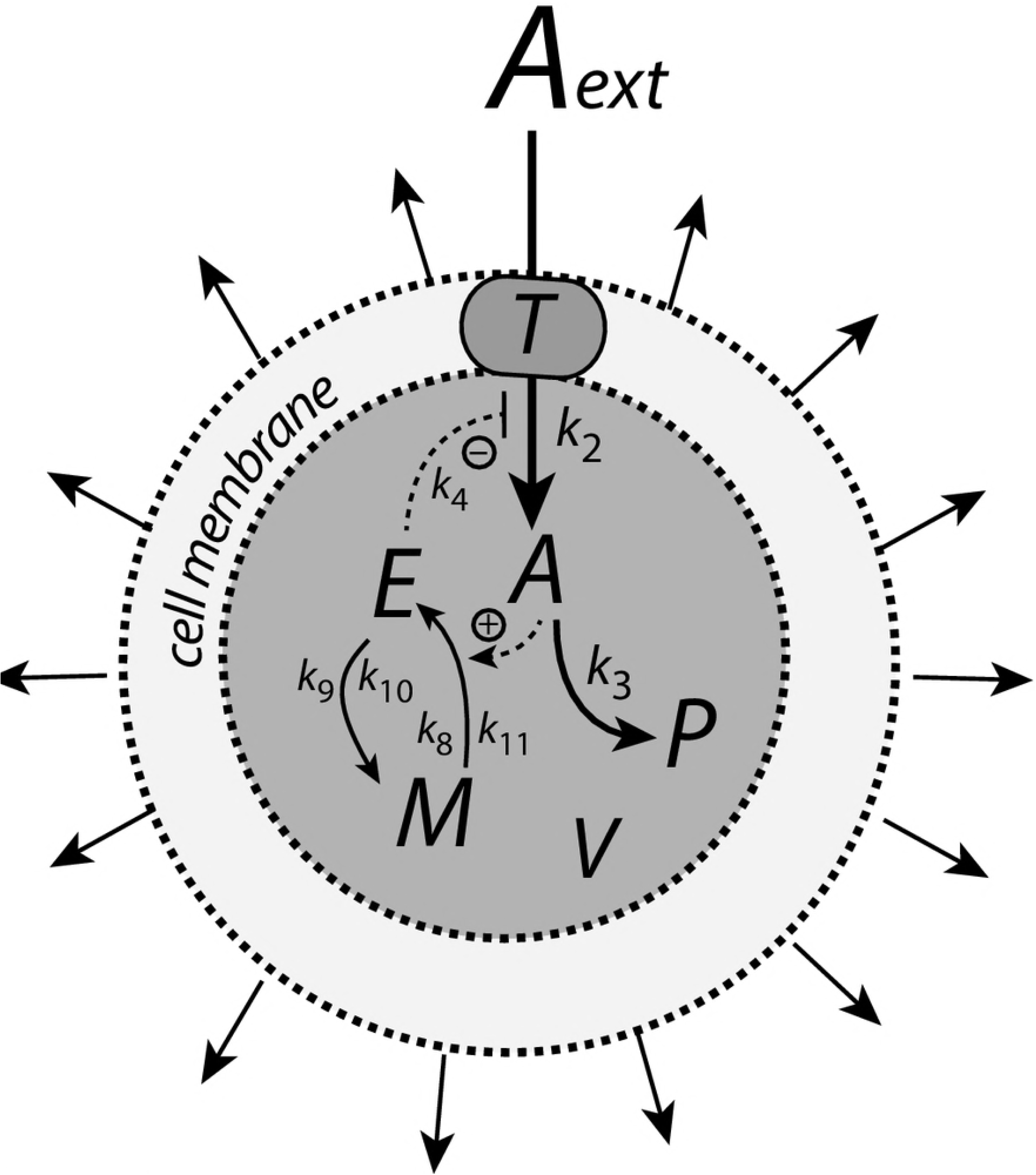
Motif 2 based controller with zero-order integral control. An increase of the compensatory flux occurs by a decrease of *E* (derepression of the compensatory flux).

The rate equations are

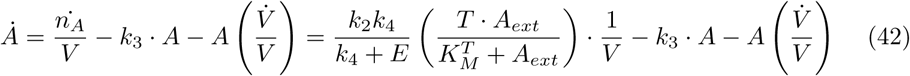

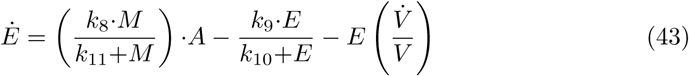

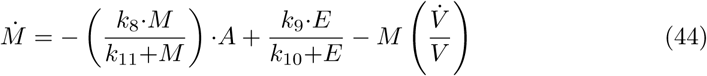

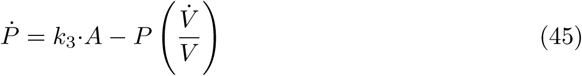

Also here we keep, for the sake of simplicity, *T* · *A_ext_/*(
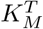
+*A_ext_*)=1. In presence of growing *V* and *k*_3_ the motif 2 zero-order controller successfully defends its theoretical set-point given by (S6 Text)

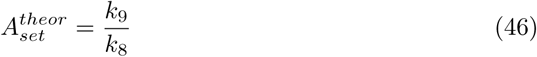

However, since an increase of the compensatory flux is based on derepression by *E* (decreasing *E*), the controller will break down when *E* ≪ *k*_4_ and *k*_4_/(*k*_4_+*E*)≈1. Neglecting the *A*·*V̇/V* term, the point when the breakdown occurs can be estimated by setting Eq. 42 to zero

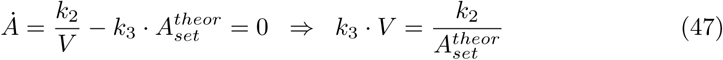

Fig. 13 shows that the motif 2 based controller is able to defend successfully against linear growth in both *V* and *k*_3_ and keeping *A* at
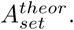
Prolonged time intervals with increasing *V* and *k*_3_ will lead to controller breakdown when the condition of Eq. 47 is met.

**Figure 13.**
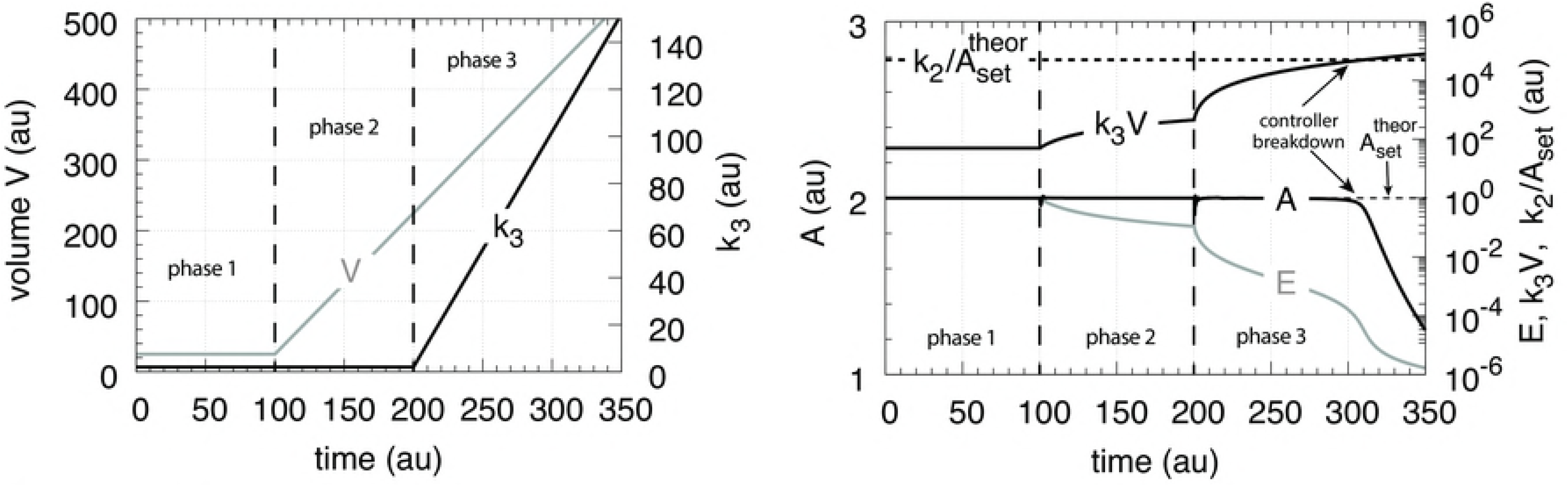
Performance of the motif 2 zero-order based controller with respect to linear increases in *V* and *k*_3_. The controller is able to defend
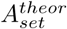
successfully, but breaks down when *k*_3_*V* reaches *k*_2_/
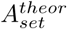
(Eq. 47). Rate parameters: *k*_2_ = 1 × 10^5^, *k*_4_ = 1 × 10^−3^, *k*_8_ = 1.0, *k*_9_ = 2.0, *k*_10_ = *k*_11_ = 1 × 10^−6^. Initial conditions: *A*_0_ =
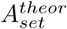
= 2.0, *E*_0_ = 1.0, *M*_0_ = 1 × 10^6^, *P*_0_ = 0.0, *V*_0_ = 25.0, *k*_3,0_ = 2.0. *V̇* =2.0 (phase 2 and phase 3), *k̇*_3_ = 1.0 (phase 3).

## Controllers with transporter-based compensatory fluxes and exponential time-dependent perturbations

Here we describe the performance of the four controller motifs (Fig. 1) with transporter-based compensatory fluxes when exposed to exponential growth, *V̇* =*k*_1_·*V*, and an exponential increase in the outflow perturbation rate parameter *k*_3_ (Fig. 14).

**Figure 14.**
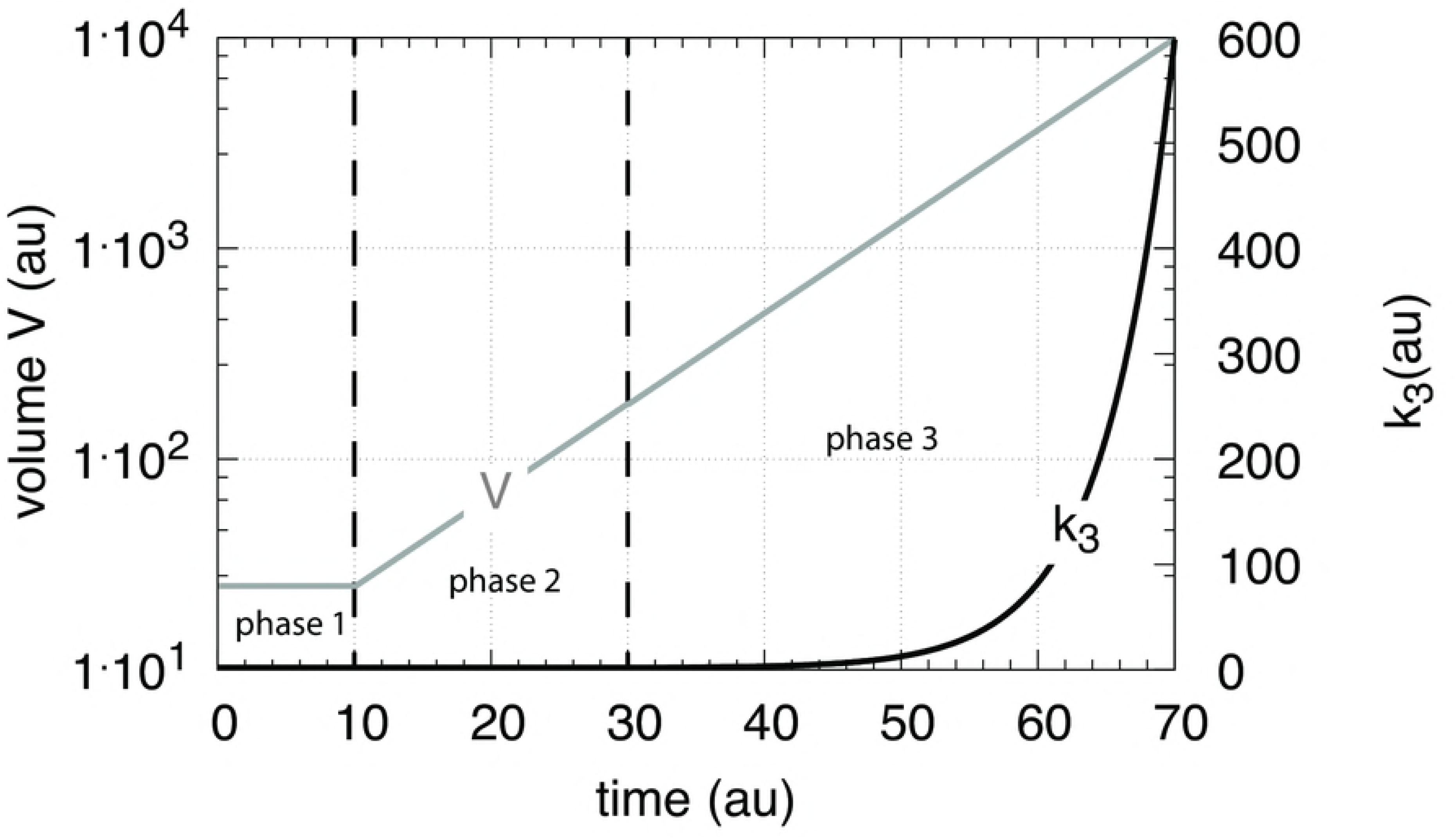
The perturbation profile with exponential growth of *V* and *k*_3_.

There are three phases the controllers are exposed to. During the first phase the controllers are at their steady states and *V* and *k*_3_ are kept constant at respectively 25.0 and 2.0. During the second phase *V* increases exponentially according to *V̇* =*k*_1_*V* (*k*_1_=0.1), while *k*_3_ is kept constant at 2.0. During phase 3, *V* continues to grow exponentially and *k*_3_ starts to increase according to

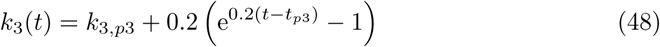

where *k*_3,*p*3_ and *t*_*p*3_ are the values of respectively *k*_3_ and time *t* at the beginning of phase 3.

Fig. 15 shows that only the motif 2 based controller with de-repression kinetics (panel d) is able to counteract both exponential increases in *V* and *k*_3_. However, due to the de-repression kinetics and due to the transporter based kinetics (see Eq. 47) the controller breaks down when the product of the perturbations, *k*_3_*V* reaches *k*_6_/
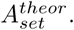
The motif 1 autocatalytical controller (panel c) shows slight constant offsets below
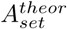
, as expected [19], both for the exponential increase of *V* and when both *V* and *k*_3_ increase exponentially. Since *E* increases with increasing perturbation strengths the controller is limited by the supply for *E* via *M* as indicated in Fig 11. Neither the motif 1 based zero-order controller (panel a) nor the antithetic controller based on motif 1 (panel b) are able to compensate for exponentially increasing perturbation strengths. They behave very similar, as already seen in Figs. 7 and 9 for linear time-dependent perturbations.

**Figure 15.**
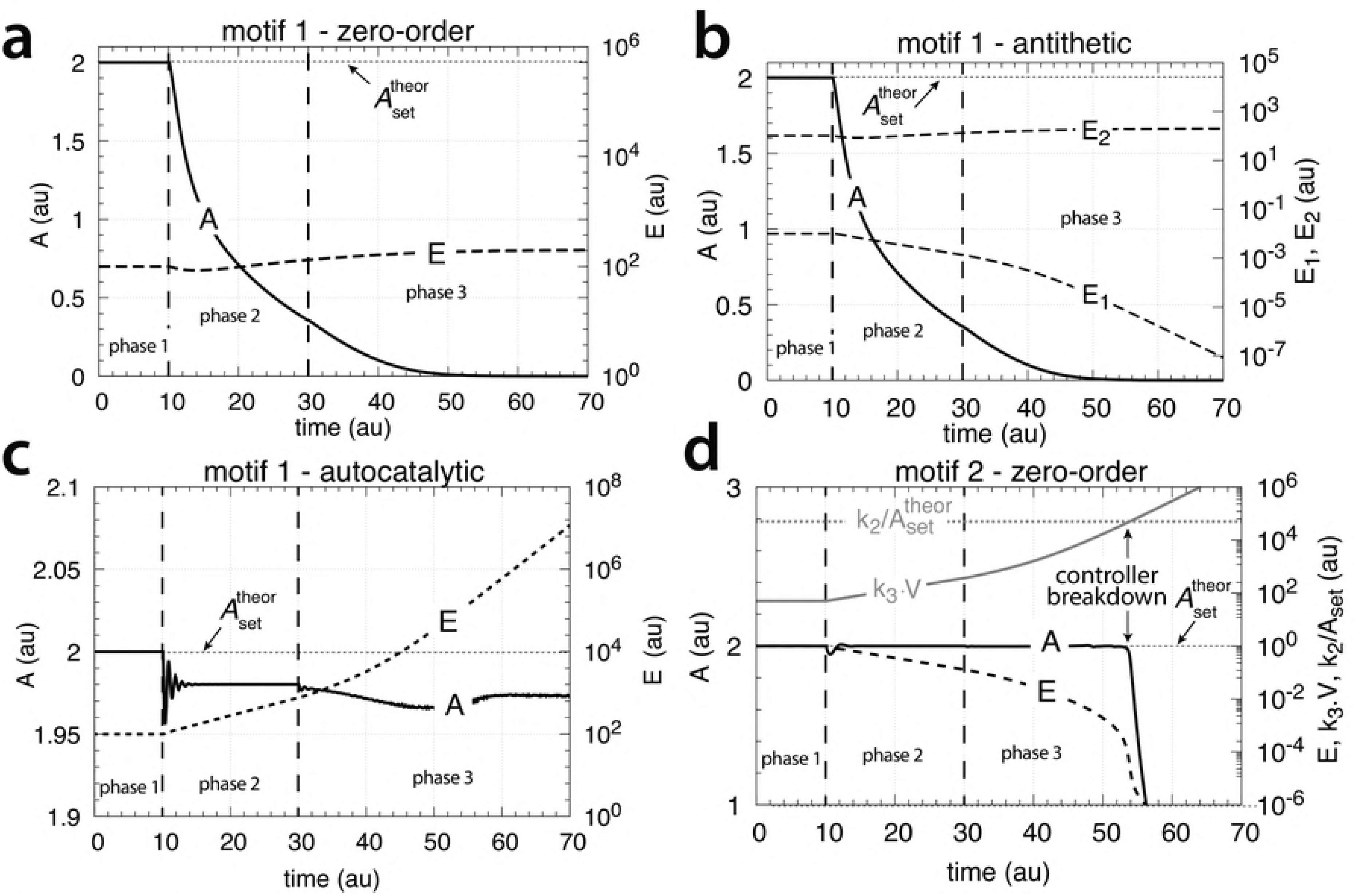
Performance of the (a) motif 1-zero-order, (b) -antithetic, (c) -autocatalytic, and (d) motif 2 zero-order controllers with transporter-based compensatory fluxes in relation to the perturbation profile of Fig. 14. For rate equations of the individual controllers, see the descriptions in the previous sections dealing with linear time-dependent perturbations. Rate parameters and initial conditions: (a) see legend of Fig. 7, (b) see Fig. 9, (c) see Fig. 11, but using *M*_0_=1×10^10^, and (d) see Fig. 13.

## Controllers with a cell-internal compensatory fluxes and linear time-dependent perturbations

We consider here the four controllers, but the compensatory fluxes are now generated from cell-internal and homogeneously distributed sources.

### Motif 1 zero-order controller

Fig. 16 shows the motif 1 zero-order controller using a cell-internal compensatory flux. The homogenously distributed compound *N* serves as a source for *A*, which is activated by *E*. Compound *M* serves as a source for *E*, while by the activation of *A*, *M* is recycled from *E*.

**Figure 16.**
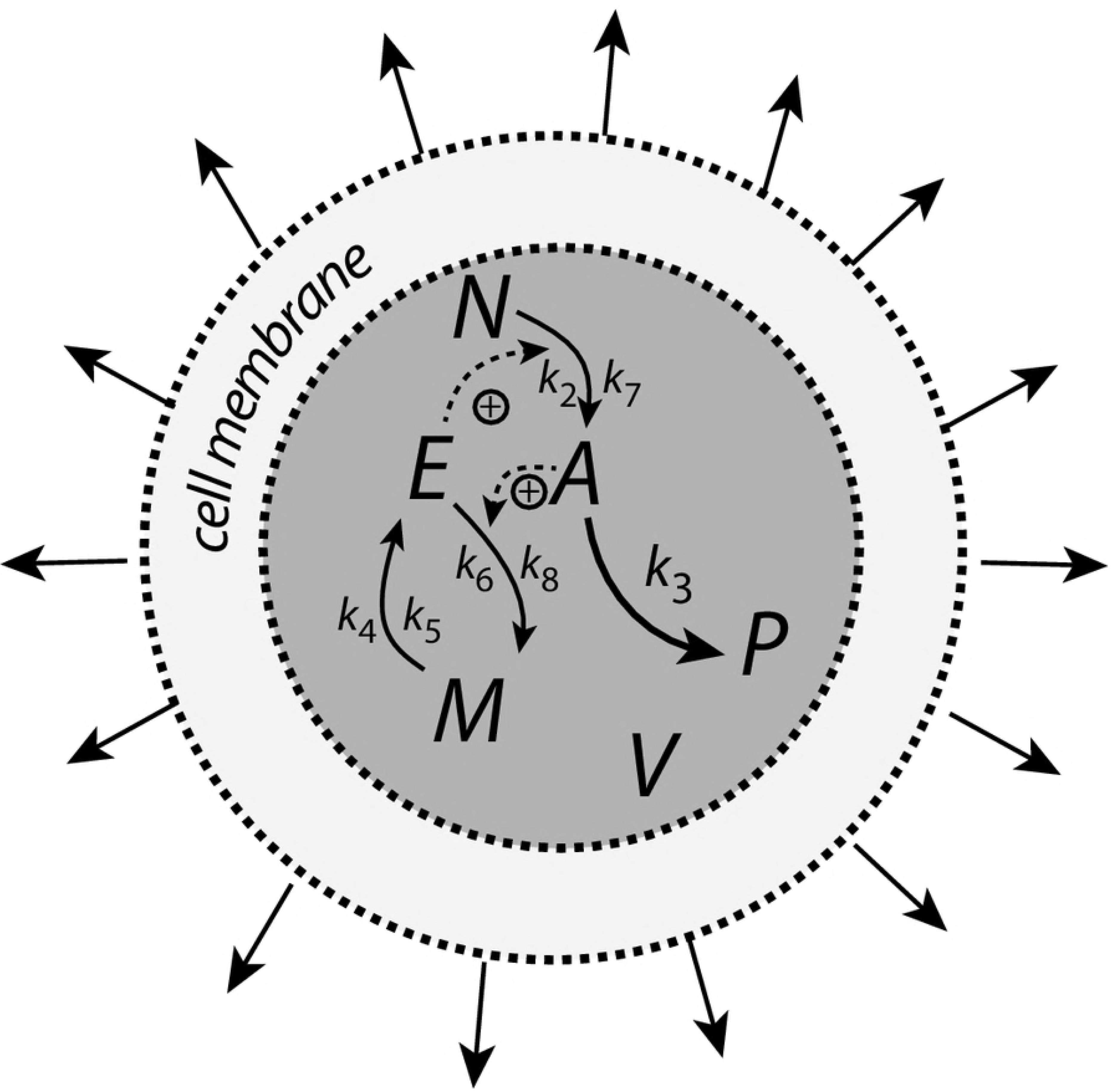
Motif 1 zero-order controller with a cell-internal compensatory flux.

The rate equations are

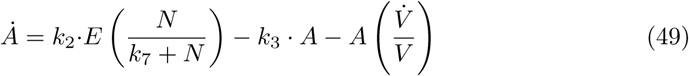

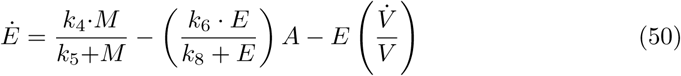

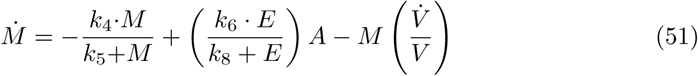

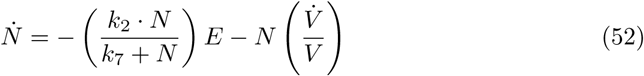

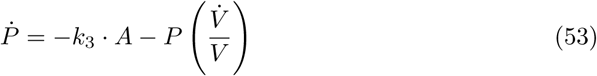

The steady state of *A* when both *V̇* and *k̇*_3_ are constant is given by the following expression (S3 Text)

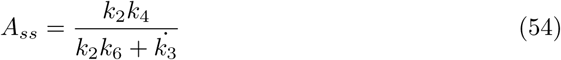

When *k̇*_3_=0 and *V̇* =constant *A_ss_* becomes
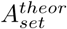
=*k*_4_/*k*_6_ and the motif 1 zero-order controller is able to compensate for a constant growth rate (Fig. 17, phases 1 and 2). However, when *k*_3_ increases linearly, *A_ss_* is below
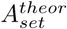
and remains constant as long as sufficient *M* and *N* are present (Fig. 17, phase 3). Thus, in comparison with a transporter-mediated compensatory fluxes, the motif 1 zero-order controller with an internally generated compensatory flux shows an improved performance by being able to compensate for a constant growth rate in the absence of other outflow perturbations in *A*.

**Figure 17.**
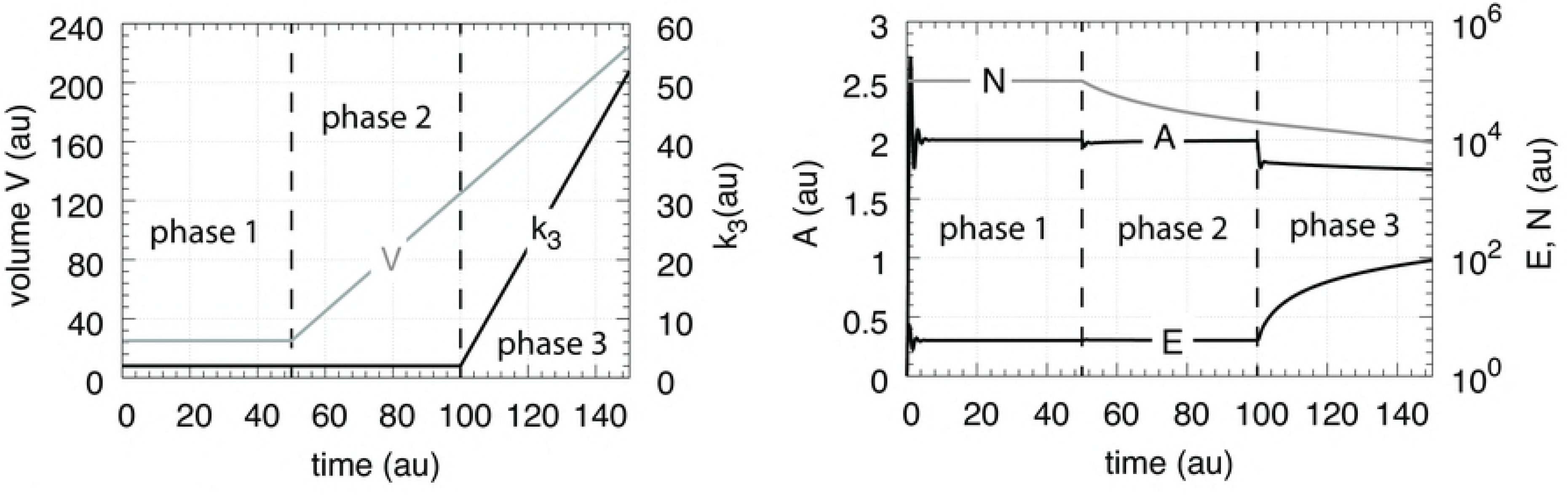
Performance of the motif 1 zero-order controller with internally generated compensatory flux (Fig. 16; Eqs. 49-53). Phase 1: constant volume *V* and constant *k*_3_. Initial volume, concentrations, and rate constants: *V*_0_=25.0, *V̇* =0.0, *A*_0_=0.0, *E*_0_=0.0, *M*_0_=4 × 10^4^, *N*_0_=1 × 10^5^, *P*_0_=0.0, *k*_2_=1.0, *k*_3_=2.0, *k̇*_3_=0.0, *k*_4_=20.0, *k*_5_=1 × 10^−6^, *k*_6_=10.0, *k*_7_=1 × 10^−6^, *k*_8_=1 × 10^−6^. The controller moves *A* to its set-point at
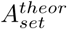
=(*k*_4_/*k*_6_)=2.0 (Eq. 54). Phase 2: rate constants remain the same as in phase 1, but *V* increases linearly with *V̇* =2.0, while *k*_3_ remains constant at *k*_3_=2.0. The controller is able to keep *A* at
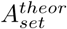
=(*k*_4_/*k*_6_)=2.0 in agreement with Eq. 54. Phase 3: *V* continues to increase with the same speed while *k*_3_ now linearly increases with *k̇*_3_=1.0. As indicated by Eq. 54 *A_ss_* leads to a constant offset below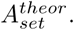

### Motif 1 antithetic controller

When the antithetic integral controller is equipped with an internally generated compensatory flux (Fig. 18) its performance towards constant growth and linearly increasing outflow perturbations *k*_3_ is significantly improved in comparison with a controller having a transporter generated compensatory flux (Fig. 9). The rate equation for *A* is now changed to

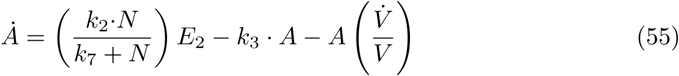

while the other rate equations (Eqs. 22-27) remain the same.

**Figure 18.**
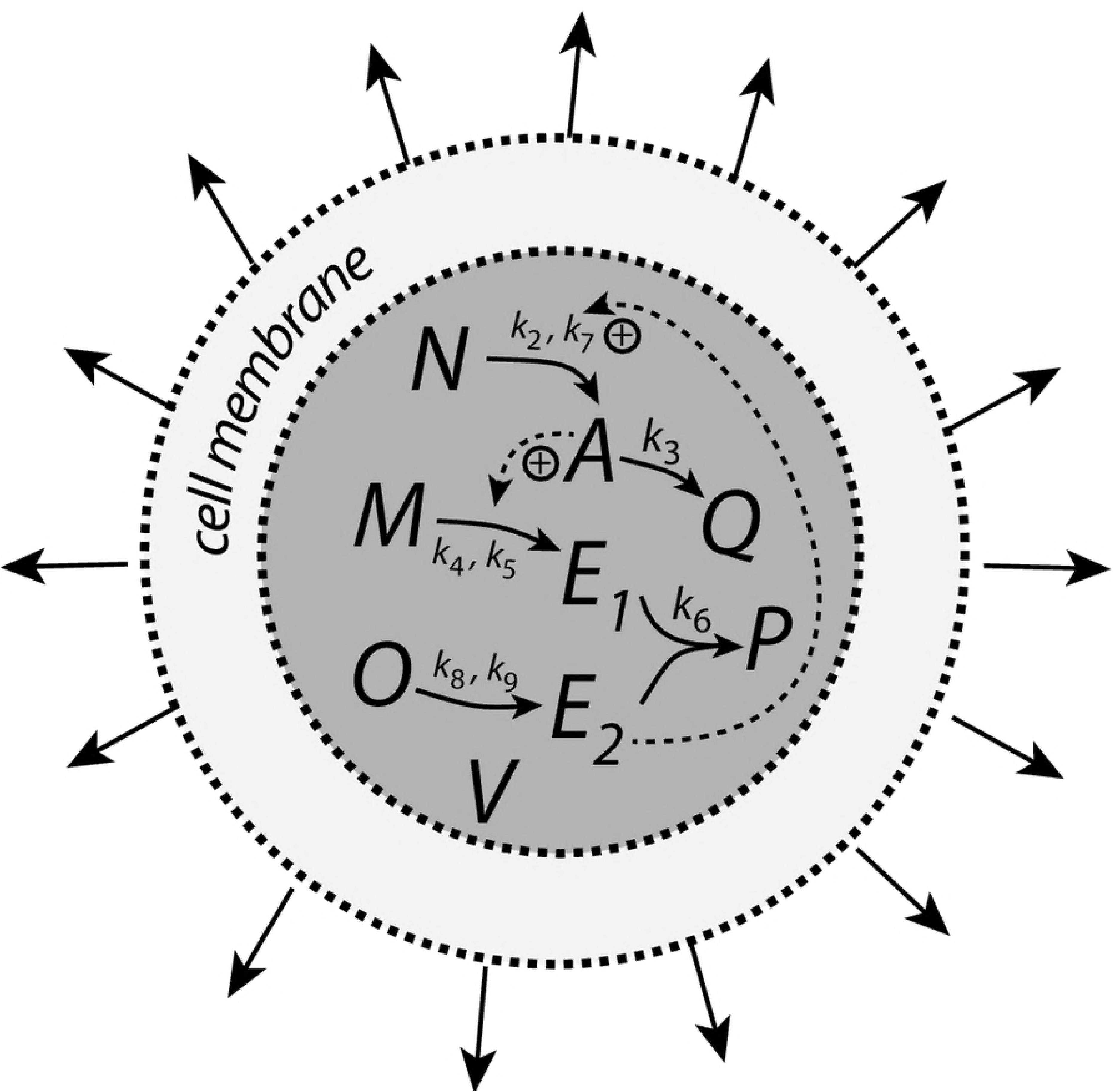
The antithetic controller with internal generated compensatory flux.

When *V̇* is constant *A_ss_* becomes (S4 Text)

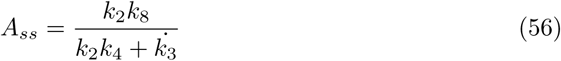

As indicated by Eq. 56 numerical results show (Fig. 19, phase 2) that the antithetic controller is now able to compensate for linear volume increases by moving *A* to
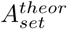
=(*k*_8_/*k*_4_). However, an offset in *A_ss_* below
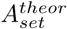
is observed when, in addition, *k*_3_ increases linearly with time, i.e., when *k*_3_ is constant.

**Figure 19.**
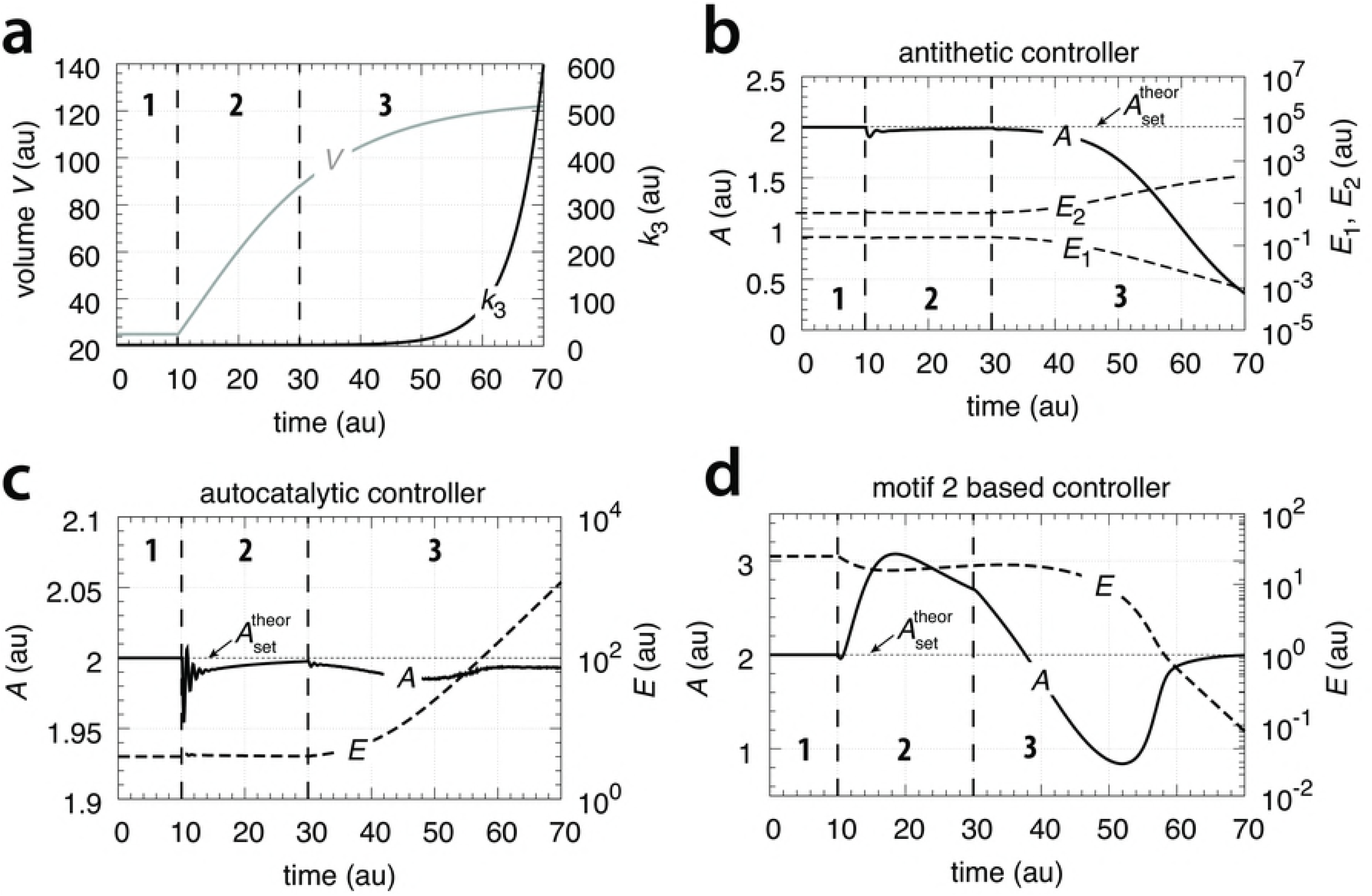
Performance of the antithetic controller when the compensatory flux is homogeneously generated within the cellular volume (Eqs. 55 and 22-27). Phase 1: constant volume *V* and constant *k*_3_. Initial concentrations and rate constant values: *V*_0_=25.0, *V̇* =0.0, *A*_0_=0.0, *E*_1,0_=0.0, *E*_2_,0=0.0, *M*_0_=2 × 10^5^, *N*_0_=1 × 10^6^, *O*_0_=2 × 10^5^, *k*_2_=1.0, *k*_3_=2.0, *k̇*_3_=0.0, *k*_4_=10.0, *k*_5_=1 × 10^−6^, *k*_6_=20.0, *k*_7_=1 × 10^−5^, *k*_8_=20.0, *k*_9_=1 × 10^−5^. The controller moves *A* to
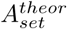
=(*k*_8_/*k*_4_)=2.0 (Eq. 56 when *k̇*_3_=0). Phase 2: rate constants remain the same as in phase 1, but *V* increases linearly with *V̇* =2.0, while *k*_3_ remains constant at *k*_3_=2.0. The controller is able to maintain *A* at
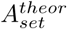
=*k*_4_/*k*_6_=2.0 in agreement with Eq. 56. Phase 3: *V* continues to increase with the same speed while *k*_3_ now linearly increases with *k̇*_3_=1.0. As indicated by Eq. 56 the controller is no longer able to keep *A* at
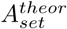
but shows a constant *A_ss_* below the theoretical set-point.

Although not explicitly shown here, during the volume increase, the mass (mole) balances described by Eqs. 30-31 are obeyed in addition to the mass balance connecting *N*, *A*, and *Q*

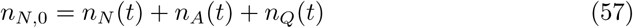

where *n*_*N*,0_ is the number of moles of initial *N* at *t* = 0 with *n*_*A*,0_=*n*_*Q*,0_=0.

### Motif 1 autocatalytic controller

Fig. 20 shows the autocatalytic controller but now with an internally generated compensatory flux. As for the motif 1 zero-order controller the compensatory flux originates from compound *N*, which is present in high concentration and forms *A* by a zero-order process with respect to *N* and activated by *E*.

**Figure 20.**
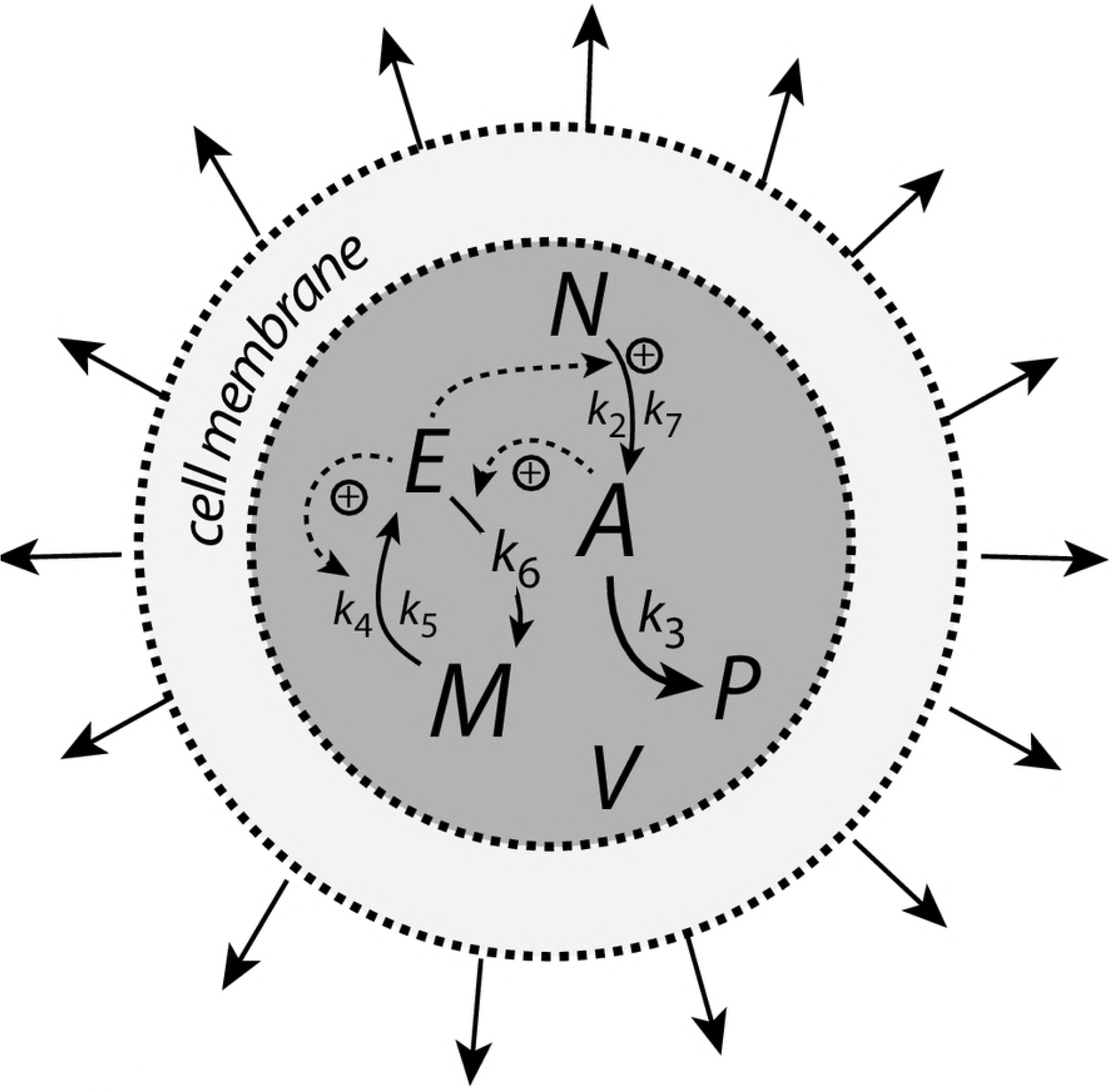
Scheme of autocatalytic controller with an internally generated compensatory flux from compound *N*. Otherwise the controller has the same structure as shown in Fig. 10.

The rate equation for the controlled variable *A* is

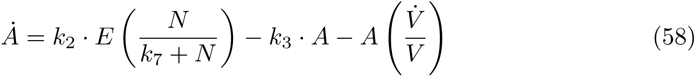

while the rate equations for *E* and *M* remain the same as Eqs. 34 and 35. Species *P* is included with the rate equation

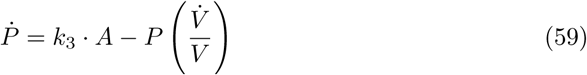

to test that the mass (mole) balance between *N*, *A*, and *P* is preserved.

The controller’s steady state in *A*, *A_ss_*, and its theoretical set-point
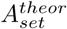
is also in case of a cell internal compensatory flux described by Eq. 38 (S5 Text). In contrast to the other controllers, even when *V̇* and *k̇*_3_ are constant the autocatalytic controller is able to move *A* to
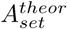
=(*k*_4_/*k*_6_) (Fig. 21).

**Figure 21.**
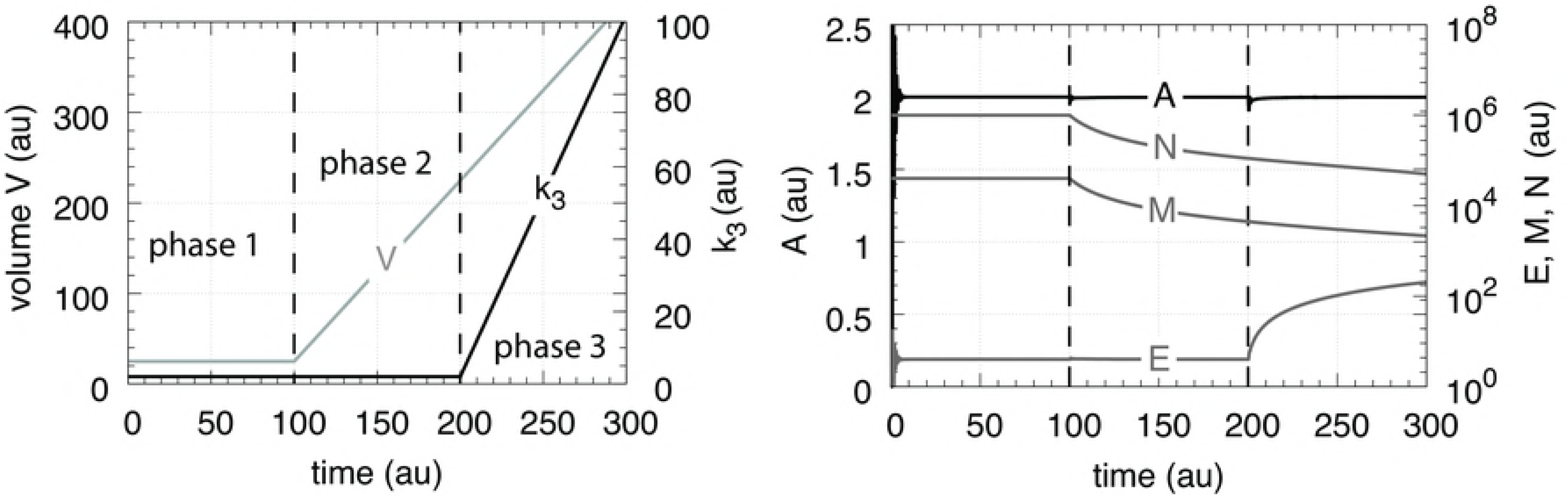
Performance of the autocatalytic controller when the compensatory flux is generated within the cellular volume (Eqs. 34-35 and 58-59). Phase 1: constant volume *V* and constant *k*_3_. Initial concentrations and rate constant values: *V*_0_=25.0, *V̇* =0.0, *A*_0_=0.0, *E*_0_=0.0, *M*_0_=4 × 10^4^, *N*_0_=1 × 10^6^, *k*_2_=1.0, *k*_3_=2.0, *k̇*_3_=0.0, *k*_4_=20.0, *k*_5_=1 × 10^−6^, *k*_6_=10.0, *k*_7_=1 × 10^−6^. The controller moves *A* to its theoretical set-point at
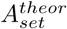
=(*k*_4_/*k*_6_)=2.0 (Eq. 37). Phase 2: rate constants remain the same as in phase 1, but *V* increases linearly with *V̇* =2.0, while *k*_3_ remains constant at *k*_3_=2.0. The controller is able to maintain *A* at
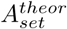
in agreement with Eq. 37. Phase 3: *V* continues to increase with the same speed while *k*_3_ now linearly increases with *k̇*_3_=1.0. As indicated by Eq. 38 the controller keeps *A* at
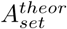
as *k*_3_ increases.

### Motif 2 zero-order controller

**Figure 22.**
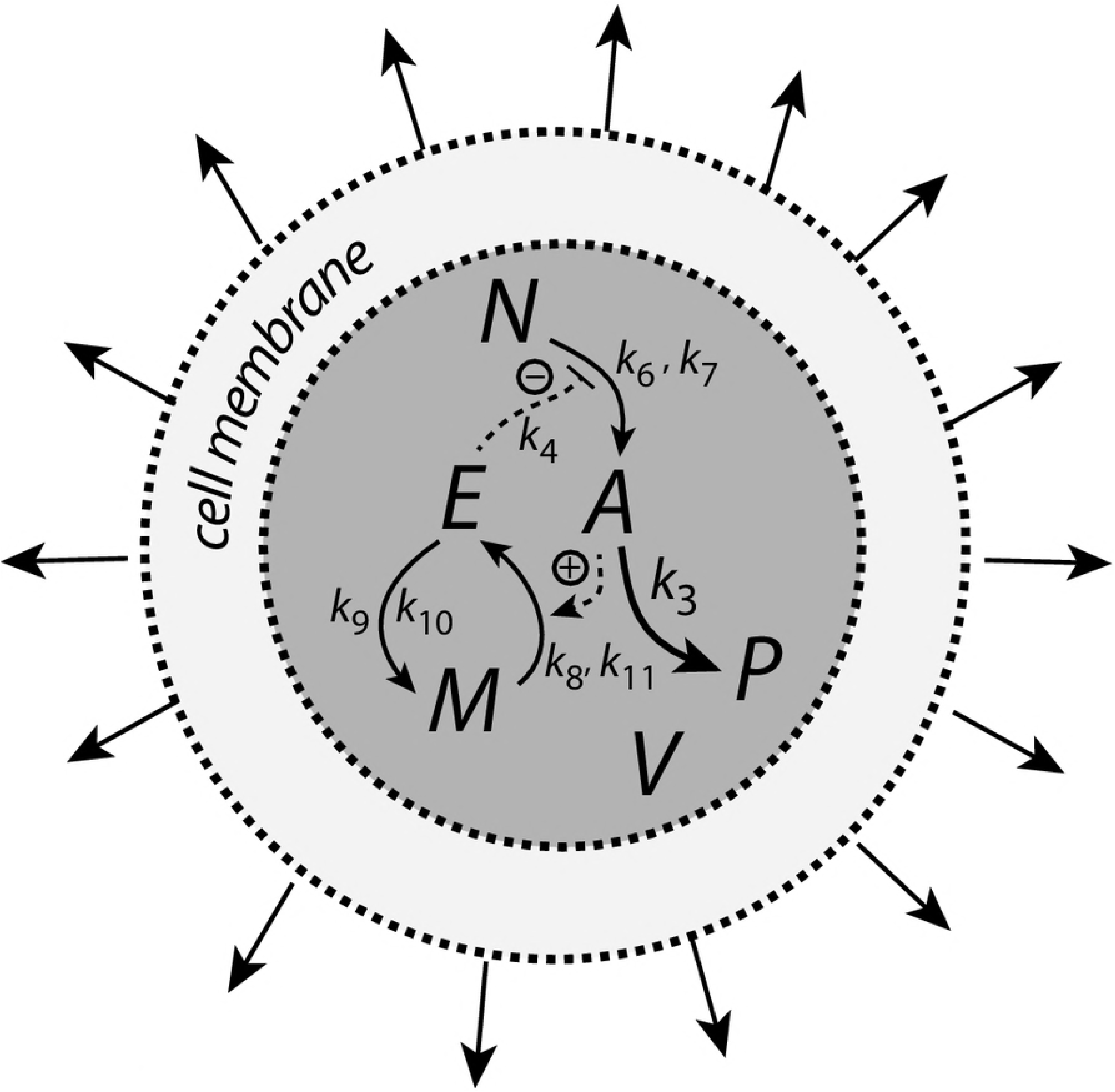
Motif 2 type controller with integral control based on zero-order kinetics and a cell-internally generated compensatory flux from compound *N*.

The rate equations for the motif 2 based controller are

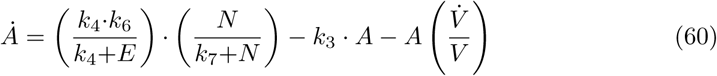

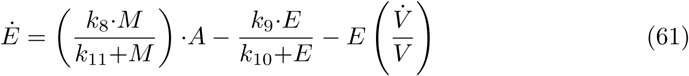

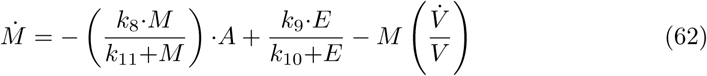

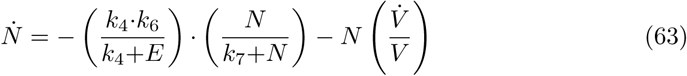

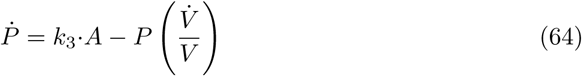

Fig. 23 shows the performance of the motif 2 controller with zero-order integral control. The controller is able to successfully defend
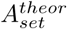
against a linear increase in *V* (phase 2) as well as against linear increase in *V* and a simultaneous linear increase in *k*_3_ (phase 3). For both cases the controller will move *A* precisely to
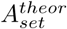
=*k*_9_/*k*_8_ without any offset (see S6 Text for details).

**Figure 23.**
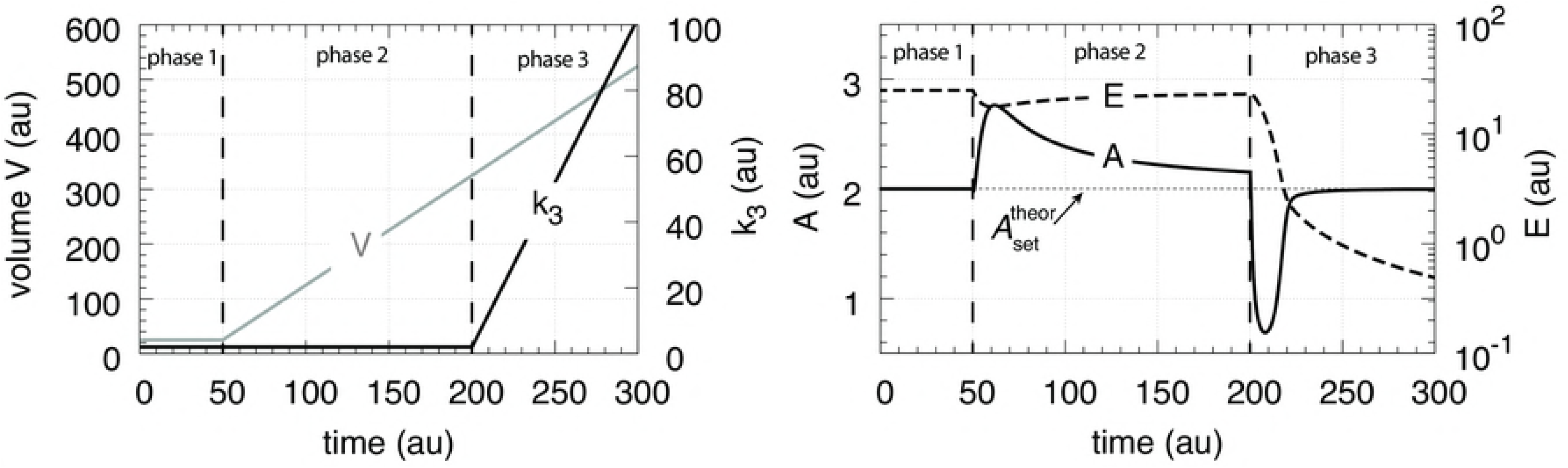
Performance of the motif 2 type of controller with zero-order based integral control. Rate constants and initial conditions: *k*_3_=2.0, *k*_4_=1 × 10^−3^, *k*_6_=1 × 10^5^, *k*_7_=1 × 10^−6^, *k*_8_=1.0, *k*_9_=2.0, *k*_10_=*k*_11_= 1 × 10^−6^, *A*_0_=2.0, *E*_0_=*V*_0_=25.0, *M*_0_=1 × 10^6^, *N*_0_=3 × 10^6^. Phase 1: *V* and *k*_3_ remain unchanged. Phase 2: *V* increases linearly with *V̇* =2.0, while *k*_3_ remains constant. Phase 3: *V* continues to increase and *k*_3_ increases linearly with *k̇*_3_=1.0.

## Controllers with cell-internal compensatory fluxes and exponential time-dependent perturbations

The controllers are exposed to the exponential perturbation profiles as shown in Fig. 14. The exponential growth of *V* is written as *V̇* = *κ* · *V*, where *κ* (>0) is a constant.

Fig. 24a shows the performance of the motif 1 zero-order controller while Fig. 24b shows the responses of the motif 1 antithetic controller. During exponential growth and constant *k*_3_ the motif 1 zero-order and the antithetic controller show slight offsets from the theoretical set-point
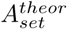
, while during phase 3 when both *V* and *k*_3_ increase exponentially, both controllers break down. Besides their different kinetic implementation of integral control both the motif 1 zero-order and the modtif 1 antithetic controller show similar responses due to analogous steady stae expressions in *A* (for details, see S3 Text and S4 Text). Fig. 24c shows the response of the autocatalytic controller, which is able to keep *A* at
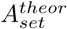
during exponential growth while *k*_3_ is kept constant. Only when *V* and *k*_3_ both increase exponentially then there is an offset from
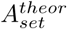
, which can be estimated analytically from the steady state condition for *A*

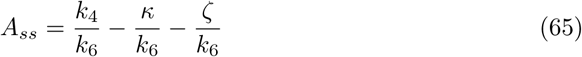

where the theoretical set-point
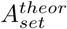
=*k*_4_/*k*_6_ and *κ* and *ζ* describe the doubling times ln 2/*κ* and ln 2/*ζ* of the exponential increases for *V* and *k*_3_, respectively (see S5 Text).

**Figure 24.**
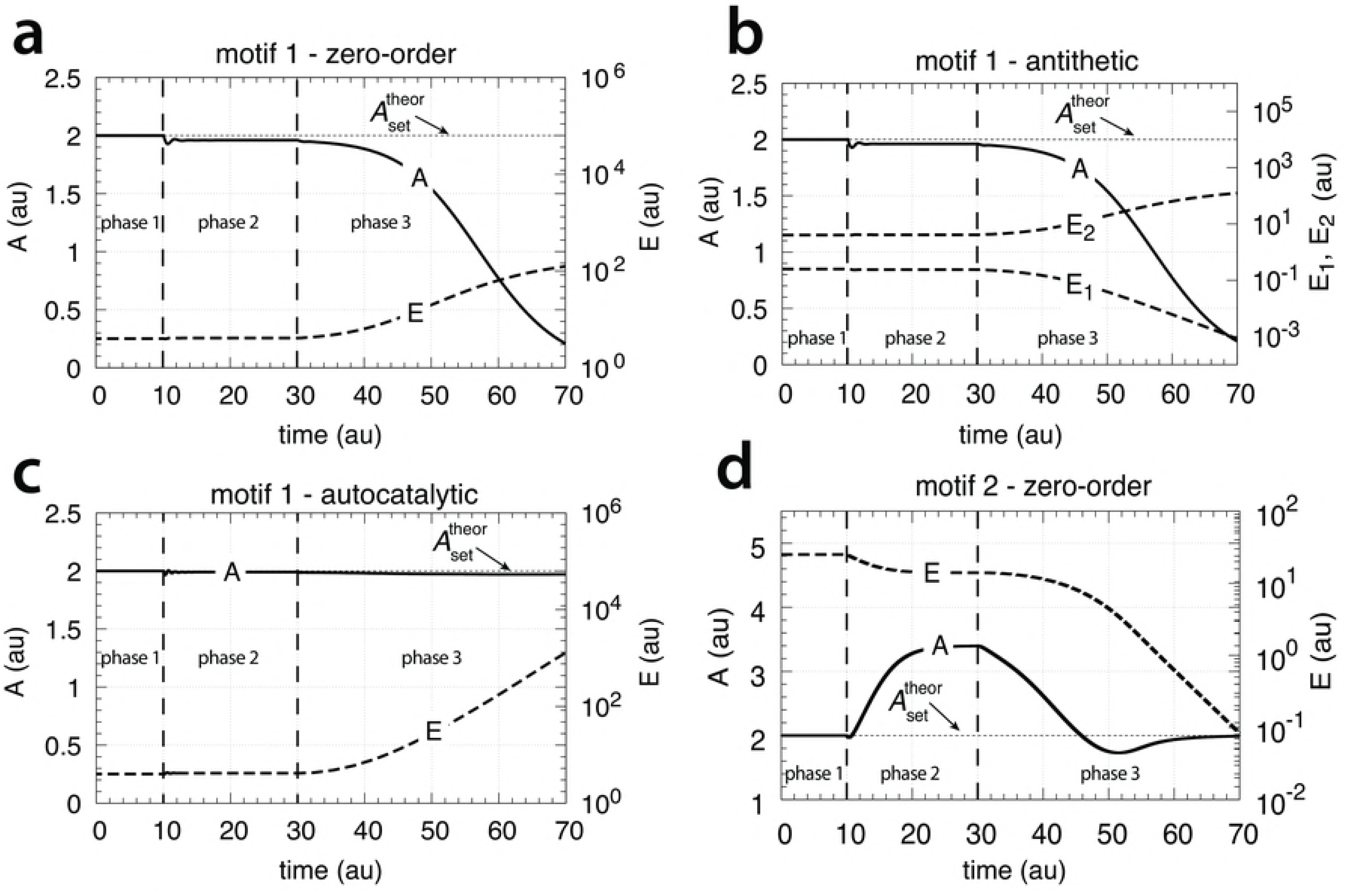
Behaviors of the motif 1 zero-order, antithetic, autocatalytic and motif 2 zero-order controllers with internal compensatory fluxes in response to an exponential increase in *V* and *k*_3_. All controllers have internal compensatory fluxes. Time/perturbation profiles of *V* and *k*_3_ are the same as in Fig. 14. (a) Behavior of the motif 1 zero-order controller. Rate constant values as in Fig. 17. Initial concentrations: *A*_0_=2.0, *E*_0_=4.0, *V*_0_=25.0, *M*_0_=4 × 10^9^, *N*_0_=1 × 10^6^. (b) Behavior of the antithetic controller. Rate constants as in Fig. 19. Initial concentrations: *A*_0_=2.0, *E*_1,0_=0.25, *E*_2,0_=4.0, *V*_0_=25.0, *M*_0_ = *N*_0_ = *O*_0_ = 1 × 10^6^, *Q*_0_ = *P*_0_=0.0. During phase 2 the controller shows a slight but constant offset below
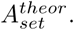
During phase 3 the controller breaks down when both *V* and *k*_3_ increase exponentially. (c) Behavior of the autocatalytic controller. Rate constants are as described in Fig. 21. Initial concentrations: *A*_0_=2.0, *E*_0_=4.0, *V*_0_=25.0, *M*_0_=4 × 10^9^, *N*_0_=1 × 10^7^. During autocatalytic growth only (phase 2) the autocatalytic controller is able to move *A_ss_* precisely to
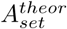
, but shows an offset from
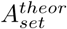
when both *k*_3_ and *V* increase exponentially). (d) Behavior of the motif 2 based controller (Eqs. 60-64). Rate constants and initial conditions as in Fig. 23. Note the significant overcompensation (offset above
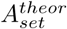)
during phase 2, but the return to
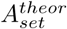
(=*k*_9_/*k*_8_) when *k*_3_ starts to grow exponentially.

The motif 2 based controller shows in phase 2 a significant overcompensation from
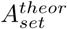
when exposed to exponential growth only. The overcompensated steady state in *A* at constant *k*_3_ and exponential growth can be expressed as

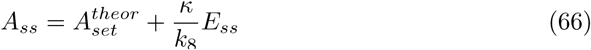

where
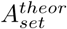
=*k*_9_/*k*_8_ and (*κ/k*_8_)*E_ss_* is the overcompensated offset (S6 Text).

The response kinetics of the motif 2 based controller is mostly determined by *k*_4_, which reflects the derepression property by *E*. For large *k*_4_ the derepression by *E* is slow and less effective.

Remarkable, when both *k*_3_ and *V* increase exponentially in phase 3 the controller is able to move *A* close to
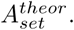
For this case *A_ss_* can be written as (S6 Text)

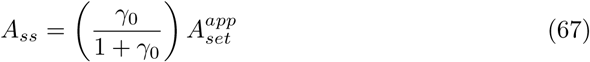

where

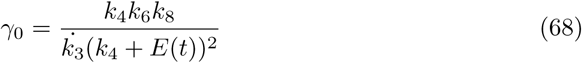

and

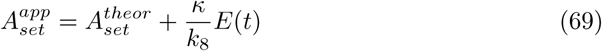

Note that during phase 3 *E* is not in a steady state, but decreases due to the controller’s derepression, while *k̇*_3_ increases exponentially. However, the derepression kinetics by *E* is faster than the exponential increase of *k̇*_3_ (Eq. 68), such that *γ*_0_ increases and
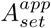
and *A_ss_* approach
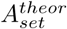
(S6 Text). A disadvantage of compensating by derepression is that during rapidly increasing *V* and *k*_3_ *E* becomes eventually so low that the negative feedback loop cannot be maintained and the controller breaks down even when sufficient *N* and *M* are available.

### Growth related to the surface to volume ratio

Here we investigate how the four controllers having cell internal compensatory fluxes perform with respect to a surface to volume ratio related growth law as found for spherical bacteria ([13, 14, 17], Eq. 2). We consider again three phases as in the previous sections, but with the difference that *V* now grows according to Eq. 2 with *η*=1 and *ξ*=0.2 (Fig. 25a). The values of *η* and *ξ* are arbitrarily chosen. The outflow perturbation, described by *k*_3_, is kept constant during phases 2 and 3, but increases during phase 3. The response behaviors of the controllers towards increasing volume (*k*_3_ is kept constant) is initially very similar to that when *V* increases linearly. However, the controllers gain control more and more control with decreasing *V̇*, provided that there is sufficient material in the cell to generate enough *E*’ (for the motif 1 controllers) or that there is still sufficient *E* left (for the motif 2 controller) to keep the negative feedback loop operating.

As an example Fig. 25 shows the behavior of the motif 1 antithetic and autocatalytic controllers and the motif 2 zero-order controller when *k*_3_ in phase 3 increases exponentially as described by Eq. 14 and compensatory fluxes are generated cell internally. The motif 1 zero-order controller’s behavior (not shown) is again very similar in comparison with the motif 1 antithetic controller.

**Figure 25.**
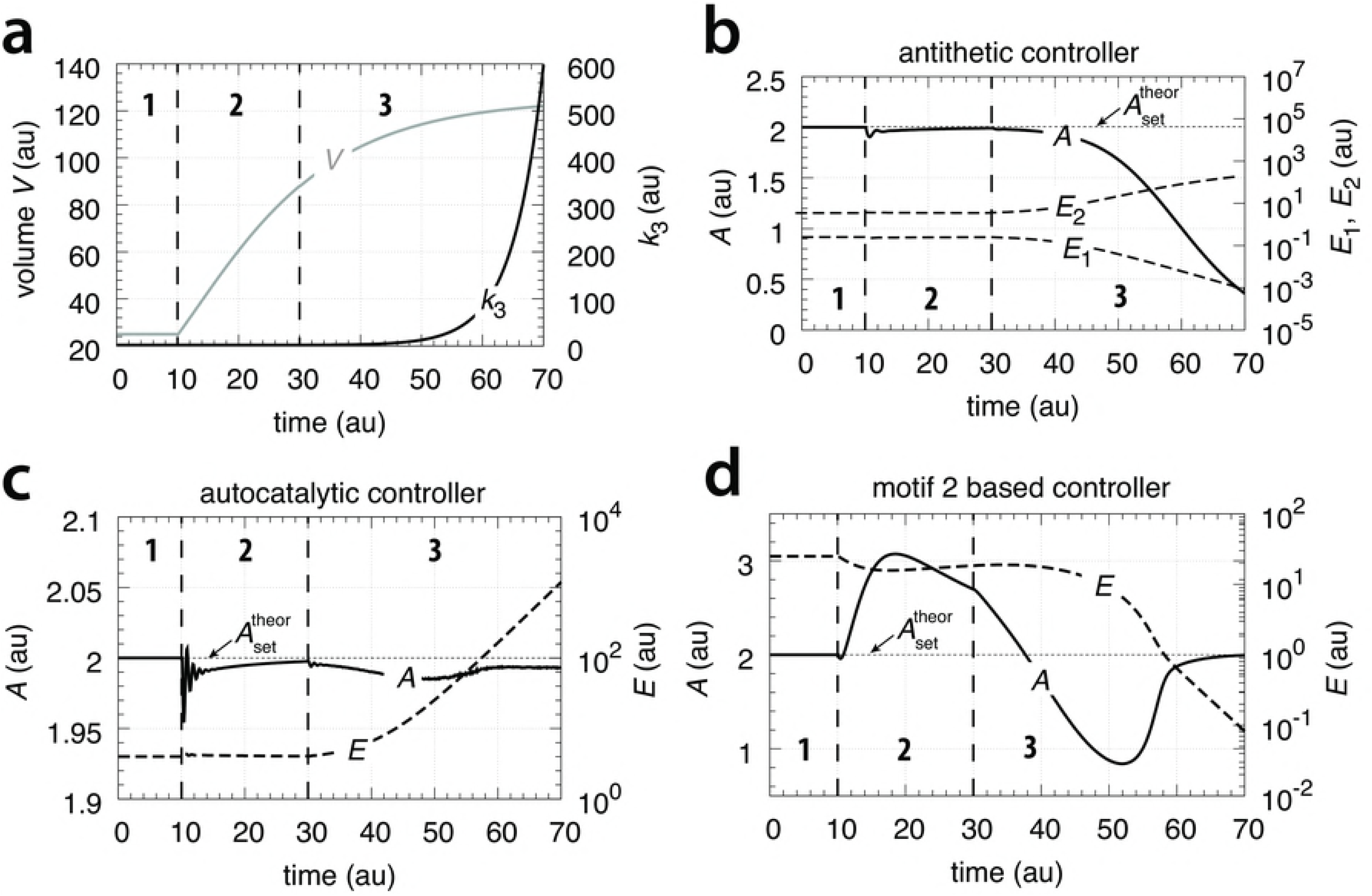
Performance of the antithetic, autocatalytic and motif 2 based controllers towards surface/volume related growth in *V* and exponentially increasing outflow perturbation *k*_3_. Rate constant values and initial conditions as in Fig. 24.

### Overview of results

Table 1 and Table 2 gives an overview of controller performances. Performances are described by the four categories *perfect adaptation*, *partial adaptation*, *over*-*adaptation*, and *breakdown*. Perfect adaptation means that the controller is able to keep *A* at
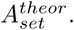
A controller with partial adaptation can maintain a constant *A* value during an applied outflow perturbation, but below
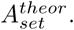
A controller showing over-adaptation keeps *A* above
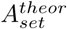
even when the perturbation should lead to a decrease in *A*. Controller breakdown means that the controller is unable to withstand the perturbation and *A* goes to zero.

**Table 1.** Performance of controllers based on internal generated compensatory fluxes

| controller type | linear $V$ only | linear $V$ and $k_3$ | exponential $V$ only | exponential $V$ and $k_3$ |
| --- | --- | --- | --- | --- |
| m1 - zero-order | perfect adaptation | partial adaptation | breakdown | breakdown |
| m1 - antithetic | perfect adaptation | partial adaptation | breakdown | breakdown |
| m1- autocatalytic | perfect adaptation | perfect adaptation | perfect adaptation | partial adaptation |
| m2 - zero-order | perfect adaptation | perfect adaptation | over-adaptation | perfect adaptation |

**Table 2.** Performance of controllers based on transporter based compensatory fluxes

| controller type | linear $V$ only | linear $V$ and $k_3$ | exponential $V$ only | exponential $V$ and $k_3$ |
| --- | --- | --- | --- | --- |
| m1 - zero-order | partial adaptation | breakdown | breakdown | breakdown |
| m1 - antithetic | partial adaptation | breakdown | breakdown | breakdown |
| m1- autocatalytic | perfect adaptation | perfect adaptation | partial adaptation | partial adaptation |
| m2 - zero-order | perfect adaptation | perfect adaptation | perfect adaptation | perfect adaptation |

Clearly, the motif 1 controllers when integral control is based on zero-order or a bimolecular (antithetic) mechanism cannot oppose an exponential volume increase or an additional exponential increase in *k*_3_. The motif 1 autocatalytic controller shows good performances with a constant offset in *A* below
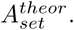
The motif 2 controller using zero-order based integral control shows best performance, is able to maintain *A* at
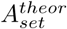
even when *V* and *k*_3_ increase exponentially. However, the drawback of the motif 2 controller is that it is based on derepression by decreasing *E* and that the controller breaks down when *E* becomes too low.

## Discussion

From Tables 1 and 2 it is seen that the motif 2 controller outperforms the other controllers. This has already been observed in a previous study [19], where time-dependent inflow and outflow perturbations for *A* at constant *V* were applied. However, a clear disadvantage of the motif 2 controller is its breakdown at low *E* value A somewhat surprising behavior of the motif 2 controller is its over-compensation whe growth increases exponentially at constant *k*_3_ (see phase 2 in Fig. 24). The over-compensation can be described analytically (Eq. 69). Its origin appears to be due to the rapid derepression kinetics. Previous results showed that the derepression kinetics are hyperbolic in nature, i.e., they can oppose growth processes with an exponentially increasing doubling time [19].

### Performance improvement by increased controller aggressiveness

Although the motif 1 zero-order and the antithetic controllers break down when expose to exponential growth and perturbations (Figs. 15 and 24), their performance can be significantly improved at constant *V̇* by increasing of what can be described as the controllers’ aggressiveness. By aggressiveness of a controller we mean loosely the controller’s response to a perturbation in terms of (mainly) quickness and precision. Increasing the aggressiveness of a controller will generally lead to a quicker controller response and an improved controller precision.

The aggressiveness of an integral controller can be varied by the controller’s gain. The gain is a factor in front of the error integral. For an ideal motif 1 zero-order integral controller (working at constant *V* and *k*_3_) *Ė* is proportional to the error *e* = (
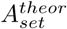
− *A*) [7], i.e.,

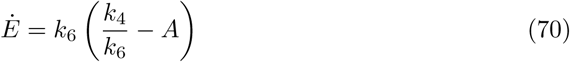

where *k*_6_ is the controller gain and *k*_4_/*k*_6_ is the controller’s theoretical set-point,
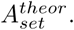
As indicated by Eq. 70 the concentration of *E* is proportional to the integrated error with respect to time. By increasing *k*_6_ and *k*_4_ such that
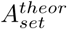
remains unchanged the gain of the controller is increased and the controller becomes more aggressive.

For constant *V̇* and *k*_3_ the steady state of *A* for the motif 1 zero-order controller is given by Eq. 20

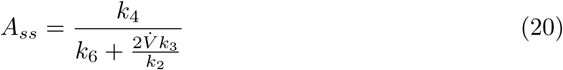

where the offset in *A_ss_* below
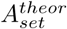
is due to the term 2*V̇k*_3_/*k*_2_. This term indicates that for increasing *V̇* and/or increasing *k*_3_ values the controller will break down and *A* will go to zero as observed in Fig. 15. There are two ways the controller’s aggressiveness can be increased. One way, as indicated above, is by increasing *k*_4_ and *k*_6_ such that
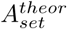
=*k*_4_/*k*_6_ is preserved with *k*_6_ becoming much larger than 2*V̇ k*_3_/*k*_2_. As a result the controller’s response kinetics become quicker and *A_ss_* moves closer to
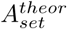
=*k*_4_/*k*_6_. The other way is to increase *k*_2_, which means to increase the activity of the transporter. This could be done by over-expressing the genes which code for the transporter.

Similar arguments apply also for the antithetic controller. Qian et al. [23] have shown that when the controller dynamics become faster than growth this leads to an improved controller performance.

Fig. 26 shows the results of increasing the aggressiveness of the motif 1 zero-order and antithetic controllers by increasing *k*_2_ from 1.0 to 1 × 10^3^. The perturbation is divided into three phases. During the first phase the volume *V* is kept constant at 25.0 and the controllers are at their set-points. In phase 2 the volume increases with a constant rate (*V̇* = 1.0). Finally, in phase 3 *V* continues to grow with *V̇* = 1.0 but *k*_2_ is increased to 1 × 10^3^. Both controllers show improved precisions, but show different kinetics in their way to reach
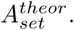

**Figure 26.**
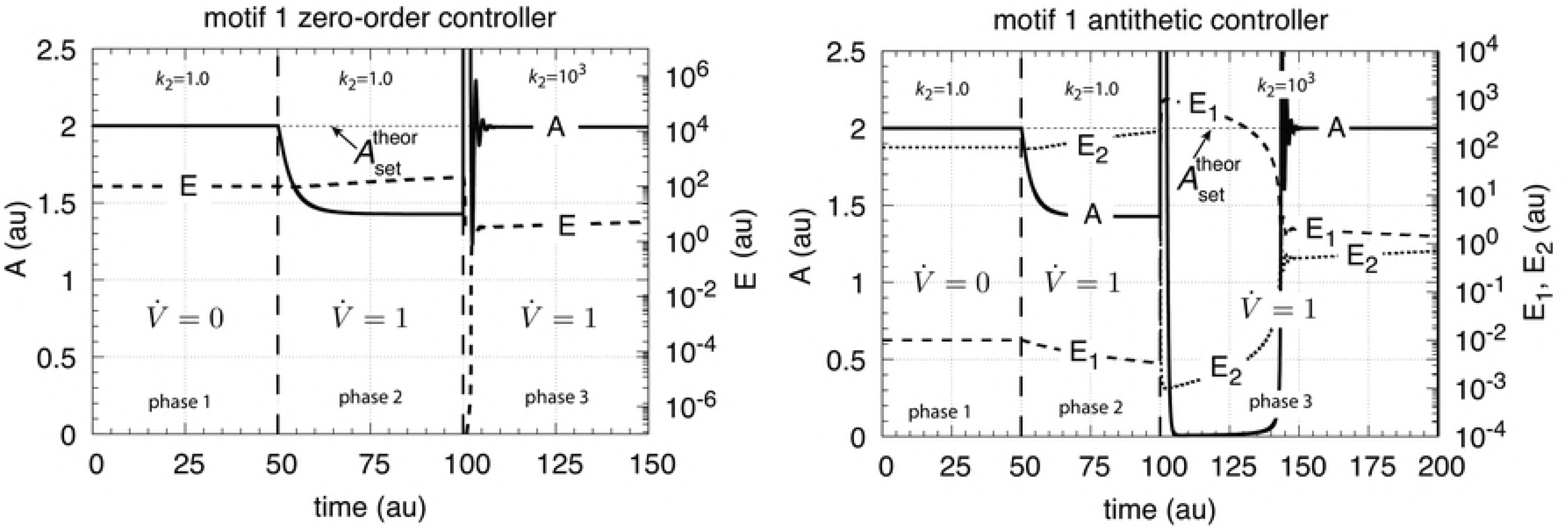
Increased transporter activity (*k*_2_ values) lead to increased aggressiveness and improved controller precision for transporter-based motif 1 zero-order controller (left panel) and motif 1 antithetic controller (right panel) during constant growth (see Figs. 6 and 8). Phase 1: controllers are at their steady state, no growth, *k*_2_=1.0. Phase 2: constant growth (*V̇* =1.0) and *k*_2_=1.0. Both controllers show an offset in *A_ss_* below
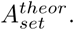
Phase 3: constant growth continues but *k*_2_ is increased to 1 × 10^3^. Both controllers show improved precision and have their *A_ss_* close to
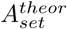
, but show different adaptation kinetics during the transition from phase 2 to phase 3. Rate parameters and initial concentrations, zero-order controller: *k*_3_=2.0, *k*_4_=20.0, *k*_5_=1 × 10^−6^, *k*_6_=10.0, *k*_7_=1 × 10^−6^, *A*_0_=2.0, *E*_0_=100.0, *V*_0_=25.0, *M*_0_=1 × 10^7^. Rate parameters and initial concentrations, antithetic controller: *k*_3_=2.0, *k*_4_=10.0, *k*_5_=1 × 10^−6^, *k*_6_=10.0, *k*_8_=20.0, *k*_9_=1 × 10^−6^, *A*_0_=2.0, *E*_1_,0=1 × 10^−2^, *E*_2_,0=1 × 10^2^, *V*_0_=25.0, *M*_0_=*O*_0_=1 × 10^8^.

Similar is the situation when the compensatory flux is internally generated. Eq. 54 shows the steady state in *A* for the motif 1 zero-order controller. Also here increasing *k*_2_ values will move *A_ss_* towards the theoretical set-point
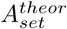
=*k*_4_/*k*_6_.

## Roles of kinetic implementations of integral control and negative feedback structures

The increased aggressiveness of the motif 1 zero-order and antithetic controllers allows them to defend their theoretical set-points as long as

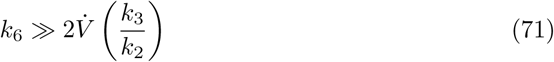

However, for exponentially increasing *V* and *V̇*
this can be achieved only for a
certain (often short) time period. The motif 1 zero-order controller will break down when Eq. 71 is no longer fulfilled. On the other hand, as shown above, the autocatalytic motif 1 controller is able to maintain a stable steady state in *A*, although with an offset from
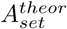
, when *V* and *k*_3_ increase exponentially. As Eq. 38 (for the transporter-based compensatory flux) indicates, any time-dependent perturbation of the type *k*_3_(*t*) = *k*_3,0_ + *a*·*t* (*a* > 0) will be successfully defended by the autocatalytic controller, because *k̇*_3_/*k*_3_ → 0 and thereby restoring the controller’s theoretical set-point. However, breakdown may occur if no sufficient supply of *E* (for example via *M*, Fig. 11) can be maintained.

Our results indicate that the type of kinetics realizing integral control (for example by autocatalysis) and the structure of the negative feedback loop (motifs 1-8, [7]) play essential roles in how a controller will perform. For example, the occurrence of autocatalytic steps/positive feedback loops in signaling and within homeostatic regulated feedback loops are becoming recognized [24–27]. As an illustration, in cortisol homeostasis ACTH signaling from the brain-pituitary system to the cortisol producing adrenals occurs by autocatalysis/positive feedback. In blood sugar homeostasis at low insulin concentrations insulin acts on its own secretion and leads to an autocatalytic production of insulin [27]. These examples indicate the importance of additional “helper kinetics” (such as autocatalysis/positive feedback) to obtain a homeostatic regulation with optimum response and precision properties.

In control engineering the *Internal Model Principle* [28–30] states that in order adapt to an environmental perturbation the controller needs to have the capability generate that type of perturbation internally. For synthetic biology, this indicates that knowledge about controller kinetics and controller structure (single or combined controller motifs) will be important for succeeding in the design and implementation of artificial regulatory units in order to oppose the dilution effect of growth or other time-dependent perturbations.

## Supporting Information

**S1 Matlab. A set of Matlab files showing the results of Figs. 7, 9, 11, 13, 15, 17, 19, 21, 23, 24, 25, and 26.**

(ZIP)

**S1 Text. Steady state of cell-internal-generated compound** *A* **without negative feedback**

(PDF)

**S2 Text. Steady state of transporter-generated compound** *A* **without negative feedback**

(PDF)

**S3 Text. Steady states and theoretical set-point for motif 1 zero-order controller**

(PDF)

**S4 Text. Steady states and theoretical set-point for motif 1 second-order (antithetic) controller**

(PDF)

**S5 Text. Steady states and theoretical set-point for motif 1 autocatalytic controller**

(PDF)

**S6 Text. Steady states and theoretical set-point for motif 2 zero-order controller**

(PDF)

## Acknowledgments

This research was financed in part by Program Area Funds from the University of Stavanger to PR and TD.

## References

1. Cannon W. Organization for Physiological Homeostatics. Physiol Rev. 1929;9:399–431.

2. Langley, LL, editor. Homeostasis. Origins of the Concept. Stroudsbourg, Pennsylvania: Dowden, Hutchinson & Ross, Inc.; 1973.

3. Bernard C. An Introduction to the Study of Experimental Medicine. English translation of the 1865 French edition by Henry Copley Greene. Dover: Macmillan & Co., Ltd; 1957.

4. Mrosovsky N. Rheostasis. The Physiology of Change. New York: Oxford University Press; 1990.

5. Risvoll GB, Thorsen K, Ruoff P, Drengstig T. Variable setpoint as a relaxing component in physiological control. Physiological Reports. 2017;5(17):e13408.

6. Ni XY, Drengstig T, Ruoff P. The control of the controller: Molecular mechanisms for robust perfect adaptation and temperature compensation. Biophys J. 2009;97(5):1244–1253.

7. Drengstig T, Jolma I, Ni X, Thorsen K, Xu X, Ruoff P. A basic set of homeostatic controller motifs. Biophys J. 2012;103(9):2000–2010.

8. Thorsen K, Agafonov O, Selstø CH, Jolma IW, Ni XY, Drengstig T, et al. Robust Concentration and Frequency Control in Oscillatory Homeostats. PLOS ONE. 2014;9(9):e107766. doi:10.1371/journal.pone.0107766.

9. Wilkie J, Johnson M, Reza K. Control Engineering. An Introductory Course. New York: Palgrave; 2002.

10. Yi TM, Huang Y, Simon MI, Doyle J. Robust perfect adaptation in bacterial chemotaxis through integral feedback control. Proc Natl Acad Sci USA. 2000;97(9):4649–53.

11. El-Samad H, Goff JP, Khammash M. Calcium homeostasis and parturient hypocalcemia: an integral feedback perspective. J Theor Biol. 2002;214(1):17–29.

12. Kitano H. Towards a theory of biological robustness. Mol Sys Biol. 2007;3:1–7.

13. von Bertalanffy L. Metabolic types and growth types. The American Naturalist. 1951;85(821):111–117.

14. Von Bertalanffy L. Quantitative laws in metabolism and growth. The Quarterly Review of Biology. 1957;32(3):217–231.

15. Smith JH. On the early growth rate of the individual fungus hypha. New Phytologist. 1924;23(2):65–78.

16. Schmalhausen I, Bordzilowskaja N. Das Wachstum niederer Organismen. I. Das Individuelle Wachstum der Bakterien und Hefe. Wilhelm Roux’Archiv für Entwicklungsmechanik der Organismen. 1930;121(4):726–754.

17. Bertalanffy L. Principles and Theory of Growth. In: Nowinski WW, editor. Fundamental Aspects of Normal and Malignant Growth. Amsterdam: Elsevier; 1960. p. 137–259.

18. Radhakrishnan K, Hindmarsh AC. Description and Use of LSODE, the Livermore Solver for Ordinary Differential Equations. NASA Reference Publication 1327, Lawrence Livermore National Laboratory Report UCRL-ID-113855. Cleveland, OH 44135-3191: National Aeronautics and Space Administration, Lewis Research Center; 1993.

19. Fjeld G, Thorsen K, Drengstig T, Ruoff P. The Performance of Homeostatic Controller Motifs Dealing with Perturbations of Rapid Growth and Depletion. J Phys Chem B. 2017;121:6097–6107.

20. Drengstig T, Ni XY, Thorsen K, Jolma IW, Ruoff P. Robust Adaptation and Homeostasis by Autocatalysis. J Phys Chem B. 2012;116:5355–5363.

21. Briat C, Gupta A, Khammash M. Antithetic integral feedback ensures robust perfect adaptation in noisy biomolecular networks. Cell Systems. 2016;2(1):15–26.

22. Thorsen K, Ruoff P, Drengstig T. Antagonistic regulation with a unique setpoint, integral and double integral action. bioRxiv. 2018;p. 248682.

23. Qian Y, Del Vecchio D. Realizing ‘integral control’in living cells: how to overcome leaky integration due to dilution? Journal of The Royal Society Interface. 2018;15(139):20170902.

24. DeAngelis DL, Post WM, Travis CC. Positive Feedback in Natural Systems. Berlin-Heidelberg: Springer-Verlag; 1986.

25. Becskei A, Séraphin B, Serrano L. Positive feedback in eukaryotic gene networks: cell differentiation by graded to binary response conversion. The EMBO Journal. 2001;20(10):2528–2535.

26. Angeli D, Ferrell JE, Sontag ED. Detection of multistability, bifurcations, and hysteresis in a large class of biological positive-feedback systems. Proc Natl Acad Sci USA. 2004;101(7):1822–1827.

27. Peters A, Conrad M, Hubold C, Schweiger U, Fischer B, Fehm HL. The principle of homeostasis in the hypothalamus-pituitary-adrenal system: new insight from positive feedback. American Journal of Physiology-Regulatory, Integrative and Comparative Physiology. 2007;293(1):R83–R98.

28. Francis BA, Wonham WM. The Internal Model Principle of Control Theory. Automatica. 1976;12(5):457–465.

29. Isidori A, Byrnes CI. Output Regulation of Nonlinear Systems. IEEE Transactions on Automatic Control. 1990;35(2):131–140.

30. Sontag ED. Adaptation and regulation with signal detection implies internal model. Systems & Control Letters. 2003;50(2):119–126.

